# LEAFY and WAPO1 jointly regulate spikelet number per spike and floret development in wheat

**DOI:** 10.1101/2023.11.02.565263

**Authors:** Francine Paraiso, Huiqiong Lin, Chengxia Li, Daniel P. Woods, Tianyu Lan, Connor Tumelty, Juan M. Debernardi, Anna Joe, Jorge Dubcovsky

**Affiliations:** Department of Plant Sciences, University of California, Davis, CA 95616, USA; Howard Hughes Medical Institute, Chevy Chase, MD 20815, USA; Institute for Plant Genetics, Heinrich Heine University, Düsseldorf 40225, Germany

**Keywords:** wheat, inflorescence development, spike, LFY, spatial transcriptomics

## Abstract

In wheat, the transition of the inflorescence meristem to a terminal spikelet (IM→TS) determines the spikelet number per spike (SNS), an important yield component. In this study, we demonstrate that the plant-specific transcription factor LEAFY (LFY) physically and genetically interacts with WHEAT ORTHOLOG OF APO1 (WAPO1) to regulate SNS and floret development. Loss-of-function mutations in either or both genes result in significant and similar reductions in SNS, as a result of a reduction in the rate of spikelet meristems formation per day. SNS is also modulated by significant genetic interactions between *LFY* and *SQUAMOSA* MADS-box genes *VRN1* and *FUL2*, which promote the IM→TS transition. Single-molecule fluorescence *in-situ* hybridization revealed a down-regulation of *LFY* and up-regulation of the *SQUAMOSA* MADS-box genes in the distal part of the developing spike during the IM→TS transition, supporting their opposite roles in the regulation of SNS in wheat. Concurrently, the overlap of *LFY* and *WAPO1* transcription domains in the developing spikelets contributes to normal floret development. Understanding the genetic network regulating SNS is a necessary first step to engineer this important agronomic trait.

**SUMMARY STATEMENT:** The plant specific transcription factor *LEAFY* plays an important role in the regulation of the number of spikelets per spike in wheat.

## INTRODUCTION

Every year, trillions of wheat spikes mature worldwide carrying the grains that provide one-fifth of the calories and proteins consumed by the human population (FAOSTAT, 2017). Therefore, increasing the maximum number of grains that can be produced by each spike can contribute to remedying the global needs for increased wheat productivity to feed a growing human population.

Wheat spikes, as other grass inflorescences, are comprised of specialized reproductive organs called spikelets, which are short indeterminate branches. Each spikelet has two proximal sterile bracts (glumes) followed by a variable number of florets. Individual florets include a lemma, which is also a bract, subtending the floral organs (one palea, two lodicules, three stamens and a pistil) (Preston *et al*., 2009, Debernardi *et al*., 2020a). The wheat inflorescence meristem (IM) produces multiple lateral spikelet meristems (SMs) in a distichous order before transitioning to a terminal spikelet (henceforth, IM→TS). The timing of this transition and the rate at which the SMs are formed determine the spikelet number per spike (SNS) and the maximum number of grains that can be formed in the spike.

The number of spikelets in a wheat spike is affected by multiple environmental conditions including drought, salt stress, heat, and reduced nutrients, all of which result in reduced SNS (Frank and Bauer, 1982, Frank *et al*., 1987, Maas and Grieve, 1990). However, differences in SNS also have a strong genetic component, with broad sense heritability ranging from *H^2^* = 0.84 in irrigated fields to *H^2^* = 0.59 in water stressed environments (Zhang *et al*., 2018). This high heritability has facilitated the identification of several wheat genes involved in the regulation of SNS. *VERNALIZATION1* (*VRN1*), *FRUITFULL2* (*FUL2*) and *FUL3*, the wheat homologs of Arabidopsis *SQUAMOSA* MADS-box genes *APETALA1* (*AP1*), *CAULIFLOWER* (*CAL*) and *FUL*, were found to be essential for spikelet development and for the regulation of the IM→TS transition (Li *et al*., 2019). Loss-of-function mutations in *vrn1* or *ful2* result in normal plants with significant increases in SNS. However, in the *vrn1 ful2* combined mutant the IM remains indeterminate and lateral spikelets are converted into tiller-like organs with vestigial floral organs. These vestigial floral organs disappear in the *vrn1 ful2 ful3* higher order mutant, in which spikelets revert to vegetative tillers subtended by leaves (Li *et al*., 2019).

Genes that regulate *VRN1* expression have been shown to affect SNS. *FT1*, the wheat homolog of Arabidopsis florigen *FLOWERING LOCUS T* (*FT*), binds directly to the *VRN1* promoter as part of a floral activation complex, and functions as a transcriptional activator (Li and Dubcovsky, 2008, Li *et al*., 2015). Mutants (or knock-down transgenic plants) of *FT1* (Lv *et al*., 2014) or its closest paralog *FT2* (Shaw *et al*., 2019) show reduced or delayed expression of *VRN1*, which is associated with significant increases in SNS. In contrast, overexpression of these genes results in a precocious IM→TS transition and spikes with very few spikelets (Lv *et al*., 2014, Shaw *et al*., 2019). Mutations in *PPD1* that reduce or delay *FT1* expression result in SNS increases (Shaw *et al*., 2013), whereas mutations in *ELF3* that result in the upregulation of *FT1* and *VRN1* expression reduce SNS (Alvarez *et al*., 2016). *bZIPC1* encodes a protein that physically interacts with FT2, and its mutants also show a large decrease in SNS (Glenn *et al*., 2023).

However, the underpinning mechanism by which the recently cloned gene *WHEAT ORTHOLOG OF APO1* (*WAPO1*) (Kuzay *et al*., 2019, Kuzay *et al*., 2022) regulates SNS has not yet been elucidated. *WAPO1* is orthologous to the *Oryza sativa* (rice) gene *ABERRANT PANICLE ORGANIZATION1* (*APO1*), and to the Arabidopsis gene *UNUSUAL FLORAL ORGANS* (*UFO*), which are both involved in floral development (Levin and Meyerowitz, 1995, Ikeda *et al*., 2007, Rieu *et al*., 2023a). In addition to floral defects, loss-of-function mutations in *WAPO1* or *APO1* result in significant reductions in SNS in wheat (Kuzay *et al*., 2022) and in the number of branches in the rice panicle (Ikeda *et al*., 2005), respectively.

In Arabidopsis, UFO physically interacts with the plant-specific transcription factor LEAFY (LFY) (Lee *et al*., 1997, Chae *et al*., 2008, Rieu *et al*., 2023b), and the interaction is conserved between the rice homologs APO1 and APO2 (Ikeda-Kawakatsu *et al*., 2012). The Arabidopsis LFY protein activates class-A MADS-box genes *AP1* (Parcy *et al*., 1998, Wagner *et al*., 1999) and *CAL* (William *et al*., 2004), which are homologous to the wheat *VRN1* and *FUL2* genes.

Since *VRN1*, *FUL2* (Li *et al*., 2019) and *WAPO1* (Kuzay *et al*., 2022) are all involved in the regulation of SNS, we investigated the role of *LFY* on wheat spike development.

In this study, we demonstrate that LFY physically interacts with WAPO1 and that plants carrying loss-of-function mutations in either or both genes result in similar floral abnormalities and similar reductions in SNS due to a reduced rate of SM formation. We also show significant genetic interactions for SNS between *LFY* and the meristem identity gene *VRN1*, which together with its closest paralog *FUL2* promote the IM→TS transition. Finally, we use single-molecule fluorescence *in-situ* hybridization (smFISH) to visualize the spatio-temporal expression profiles of these genes and other floral genes during spike development. These studies reveal a ten-fold increase in the ratio between the *SQUAMOSA* MADS-box genes (*VRN1 + FUL2*) and *LFY* in the distal part of the spike at the time of the IM→TS transition, supporting the opposite role of these genes in the regulation of SNS.

## RESULTS

### Induced loss-of-function mutations in *LFY* reduce SNS and alter floral morphology

Using our sequenced Kronos mutant population (Krasileva *et al*., 2017), we selected truncation mutations K2613 for *LFY-A* (henceforth *lfy*-*A*) and K350 for *LFY-B* (henceforth *lfy-B*). The *lfy*-*A* mutant has a G>A change in the acceptor splice site of the second intron, which results in mis-splicing of the third exon, a shift in the reading frame, and a premature stop codon that eliminates 121 amino acids (31% of the total protein, Fig. 1A). The *lfy-B* mutant has a premature stop codon at position 249 (Q249*) that truncates 37% of the protein. The eliminated amino acids in the two wheat mutants include the highly conserved LFY DNA binding domain, suggesting that the truncated proteins can no longer bind their target DNAs and, therefore, are most likely not functional (Maizel *et al*., 2005, Rieu *et al*., 2023b) (Fig. 1A). Primers used to track these mutations are described in data S1. The mutants were backcrossed to Kronos to reduce background mutations, and intercrossed with each other to select sister lines homozygous for the different mutation combinations, including the wildtype (WT), *lfy-A*, *lfy-B*, and the *lfy-A lfy-B* combined mutant, designated hereafter as *lfy*.

**Fig. 1.**
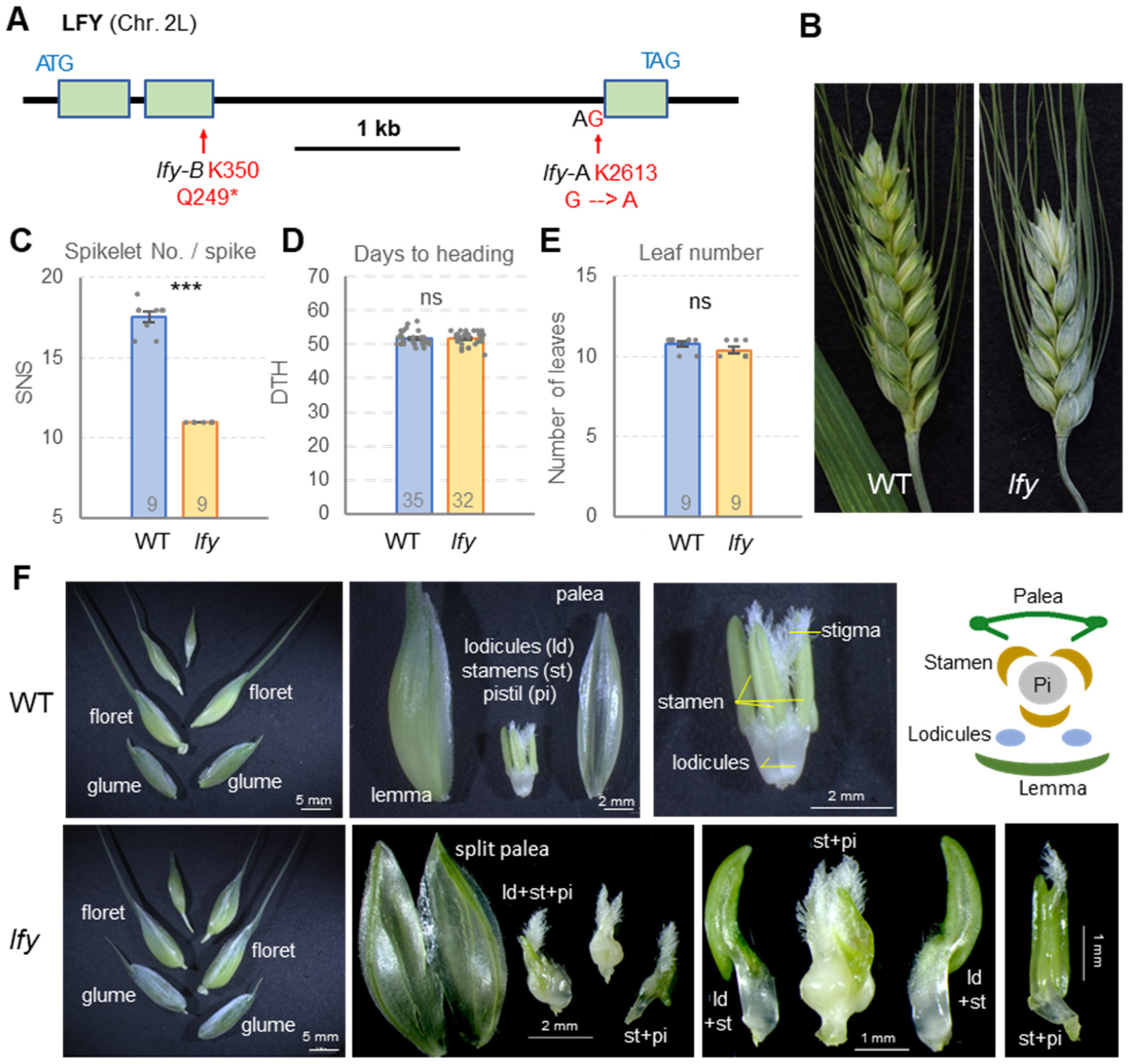
Characterization of *LFY* loss-of-function mutants. (**A**) *LFY* gene structure and selected mutations. (**B**) Representative spikes of wildtype and *lfy* mutant. Comparison between WT and *lfy* for (**C**) spikelet number per spike, (**D**) days to heading, and (**E**) leaf number. Numbers inside bars indicate biological replicates. (**F**) Top: wildtype spikelet picture and schema. Bottom: floral abnormalities in *lfy*. Raw data in data S2.

Comparisons between the homozygous sister lines in a growth chamber, revealed a highly significant decrease (37%, *P<*0.001) in SNS in the combined *lfy* mutant relative to the wildtype (WT, Fig. 1B-C, data S2). Smaller but still significant decreases in SNS were detected for the single *lfy-A* (12%) and *lfy-B* (8%) mutants (Fig. SA1, data S3), which indicates that modification of *LFY* gene dosage can be used to fine tune SNS in wheat. No significant differences in heading time or leaf number were detected between the combined *lfy* mutant and the WT (Fig. 1D-E, data S2), suggesting a limited effect of *LFY* on the timing of the transition between the vegetative and reproductive meristems.

In addition to its effects on SNS, *lfy* showed severe alterations in floral organs (Fig. 1F). We quantified the frequency of the defects in 27 first and 27 second florets from spikelets located in the basal, central and distal part of the spike (Fig. S1B-C, data S2). The glumes and lemmas developed normally, but 18.5% of the paleas were bifurcated (Fig. 1F, data S2). Eighty-one percent of the paleas were fused with either lodicules or stamens (data S2). Lodicules were also fused to stamens or membranous structures. The average number of normal stamens was reduced to 1.4 (data S2), and one fifth of the florets showed additional abnormal stamens and fusions with lodicules, membranous structures or pistils. Only 9% of the florets showed single pistils (primarily those with three normal anthers) and the rest showed more than one pistil and frequent homeotic conversions between stamens and pistils (Fig. 1F and S1, data S2).

### Overexpression of *LFY* partially rescues the reduced SNS phenotype of *lfy*

To test if *LFY* function was sufficient to rescue the mutant phenotypes, we generated transgenic plants expressing the *LFY-A1* coding region fused to a C-terminal HA tag and driven by the constitutive maize *UBIQUITIN* promoter. Transgenic lines for the five independent *UBI:LFY- HA* events, all showed significantly higher *LFY* transcript levels in the leaves than non-transgenic sister lines and wildtype Kronos, which showed no expression of endogenous *LFY* in this tissue (Fig. S2A, data S4). Among 14 dissected florets we observed missing or fused lodicules in 21%, fused stamen filaments in 43% and pistils with extra stigmas in 36% (Fig. S2B-C). Floral organ defects were less frequent and less severe than in *lfy*, which explains the higher fertility of the *UBI:LFY-HA* plants (23 ± 10 grains/plant) relative to *lfy* (2.7 ± 0.7 grains/plant), but its reduced fertility relative to the wildtype (94 ± 25 grains/plant, *P*=0.019, data S5). These results indicate that the ectopic expression of *LFY* is associated with negative pleiotropic effects on floral organ development and fertility.

We then crossed the *UBI:LFY-HA* transgenic plant #4 with *lfy*. In the progeny, we selected sister lines homozygous for combined *lfy* mutations or for wildtype alleles (WT), each with or without the transgenes. Among the plants without the transgene, the combined *lfy* mutants showed reduced SNS (6.6 spikelets, *P<*0.001), as in previous experiments. In the presence of the wildtype *LFY* alleles, transgenic plants showed 1.1 more spikelets per spike than non-transgenic controls (*P*=0.0045, Fig. 2, data S5). The effect was larger in *lfy*, where transgenic plants showed 4 more spikelets per spike than the controls (*P<*0.001, Fig. 2).

**Fig. 2.**
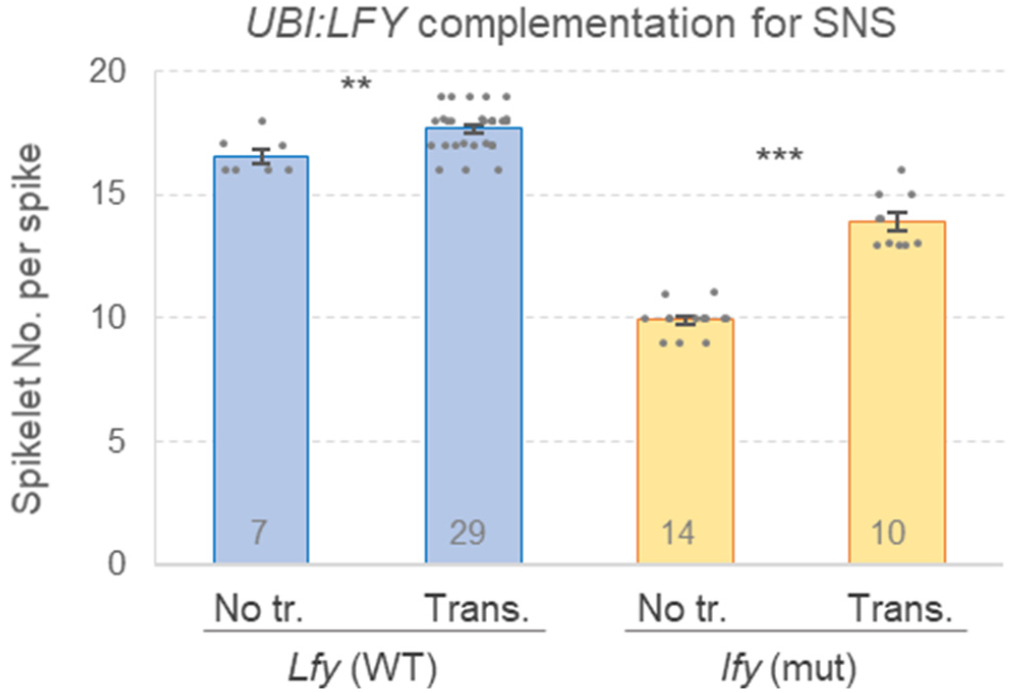
Effect of *UBI:LFY-HA* on spikelet number per spike (SNS). Effect of the *UBI:LFY- HA* transgene on SNS in wildtype and *lfy*. Numbers within bars indicate biological replicates. *P* values correspond to two-tail *t-*tests. ns = not significant, **=*P*<0.01, and ***=*P*<0.001. Raw data in data S5.

### Wheat LFY and WAPO1 show physical and genetic interactions

Since *LFY* and *WAPO1* mutants were both associated with similar reductions in SNS (Kuzay *et al*., 2022), and their homologous proteins interact with each other in Arabidopsis (Chae *et al*., 2008) and rice (Ikeda-Kawakatsu *et al*., 2012), we tested the ability of LFY and WAPO1 proteins to interact physically with each other in wheat. We used co-immunoprecipitation (Co-IP) to test the interaction between LFY-A and two WAPO-A1 natural alleles that differ in the presence of a cysteine or a phenylalanine at position 47 to test if this polymorphism affects the interaction (Kronos carries the 47C allele). The *WAPO-A1-*47F allele was previously associated with higher SNS than the *WAPO-A1-*47C allele (Kuzay *et al*., 2022). We co-transformed wheat leaf protoplasts with *UBI:LFY-HA* combined with either *UBI:WAPO1-47C-MYC* or *UBI:WAPO1-47F-MYC*. After immunoprecipitation with anti-MYC beads, we detected LFY-HA using an anti-HA antibody in both the WAPO1-47C-MYC and WAPO1-47F-MYC precipitates (Fig. 3A). These results indicate that LFY can interact with both WAPO-A1 variants in wheat.

**Fig. 3.**
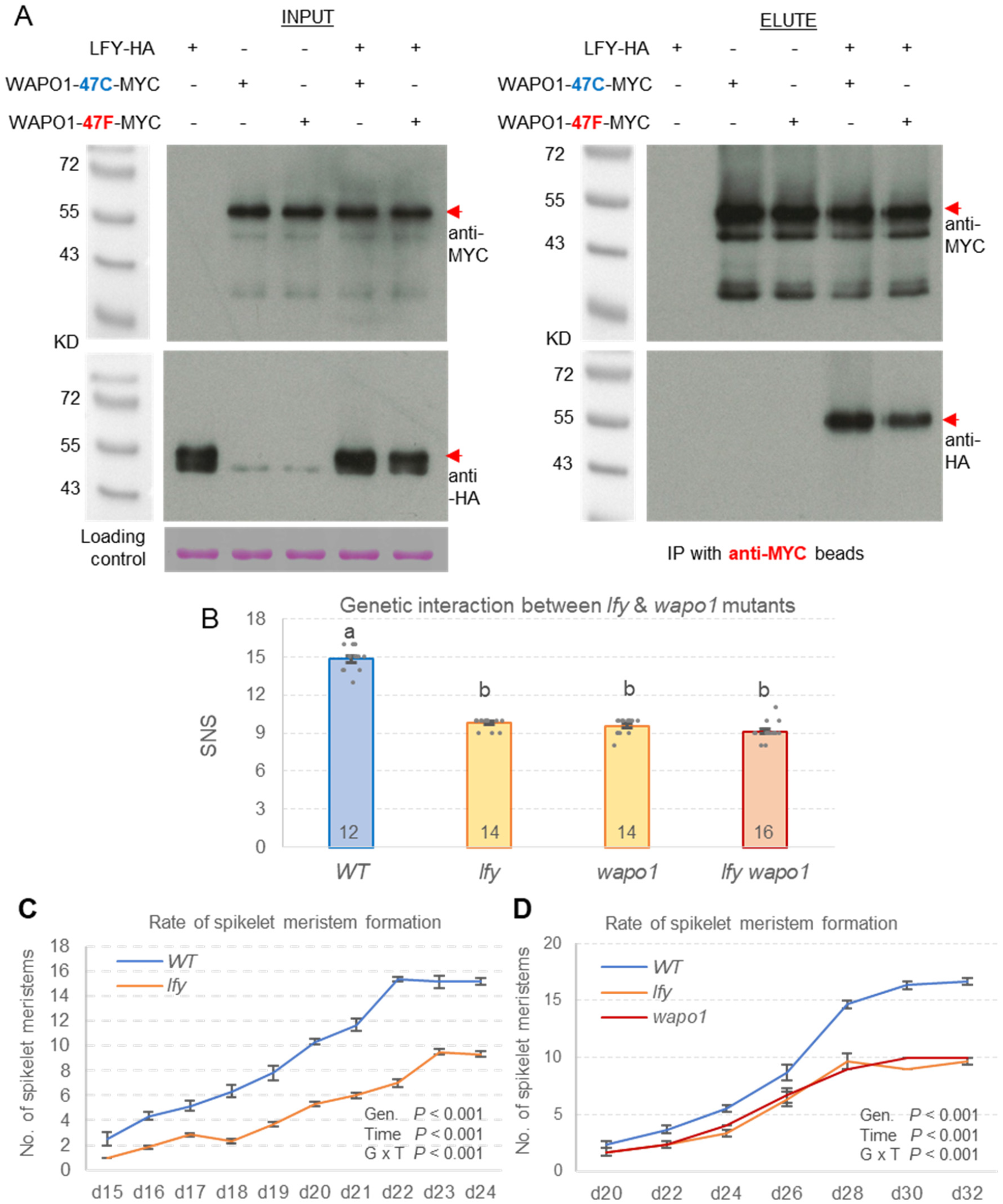
Physical and genetic interactions between WAPO1 and LFY. (**A**) Interaction between LFY and WAPO1 alleles 47F-47C in Kronos leaf-protoplasts by co-immunoprecipitation. (**B**) Genetic interaction between *lfy* and *wapo1*. Different letters indicate significant differences in Tukey tests (*P*<0.05), and numbers inside bars biological replications. (**C-D**) Changes in the number of SM with time (d=days from germination) (**C**) Wildtype and *lfy* mutant grown under long days (n=6) (**D**) Wildtype, *lfy*, and *wapo1* mutants grown 8d under short day and then long days (n=3). *P* values correspond to a repeated measures analysis. Error bars are SEM. Raw data in data S6.

To test if the physical interaction between LFY and WAPO1 was reflected in a genetic interaction for SNS, we intercrossed *lfy* with a loss-of-function *wapo1* mutant containing early truncation mutations in both *WAPO-A1* and *WAPO-B1* (Kuzay *et al*., 2022). Due to the reduced fertility of the homozygous *lfy* mutants (data S5), we first selected lines homozygous for the *lfy- A* mutant allele and heterozygous for *lfy-B* among F2 and F3 progenies. We then screened a large F4 segregating population and selected the four homozygous classes (WT, *lfy*, *wapo1*, and combined *lfy wapo1*) using molecular markers (primers in data S1). Plants homozygous for mutations in both homeologs of *LFY* (*lfy*) or *WAPO1* (*wapo1*) showed large and similar reductions in SNS relative to the wildtype (34% and 35% reduction, respectively, Fig. 3B).

Interestingly, the combined *lfy wapo1* mutant showed a reduction of 38%, relative to wildtype which was not significantly different from the reductions observed in the single mutants (Fig. 3B, data S6). The genetic epistatic interaction for SNS was highly significant in a factorial ANOVA, and the contrasts for the simple effects showed no-significant differences in SNS for *LFY* or *WAPO1* in the presence of the mutant allele of the other gene (data S6). These results indicate a reciprocal recessive epistatic interaction between these two genes, and that LFY and WAPO1 need each other to regulate SNS.

To test if the *lfy* reduction in SNS was due to a premature IM→TS transition or a reduced rate of SM formation, we dissected developing spikes and recorded the variation in SM number per day (sm/d). The first experiment (long days), showed a similar IM→TS transition time but a significantly faster rate of spikelet formation in the wildtype (1.83 sm/d) than in *lfy* (0.86 sm/d) (Fig. 3C, data S6). In the second experiment, we grew the seedlings for 8 days under short days and then transferred them to long days to synchronize the reproductive transition. In this experiment the rate of spikelet meristem formation in the wildtype (1.40 sm/d) was also faster than in the *lfy* (0.73 sm/d) and *wapo1* (0.83 sm/d, Fig. 3D). In both experiments, different rates of SM formation were observed from the earliest stages of spike development (Fig. 3C-D).

Repeated measures analyses revealed highly significant differences between mutant and wildtype genotypes, time points, and genotype x time interactions, which indicates a differential response in time (data S6). There was no significant difference in the rate of SM formation when comparing the *lfy* and *wapo1* mutants alone (data S6).

### *LFY* and *WAPO1* show dynamic expression profiles during wheat spike development

A previous RNA-seq study including different tissues at different developmental stages in Chinese Spring (CS), detected *LFY* transcripts in developing spikes and elongating stems (Choulet *et al*., 2014)(Fig. S3A). A separate RNA-seq study including five spike developmental stages in tetraploid wheat Kronos (VanGessel *et al*., 2022), showed transcripts for both *LFY* and *WAPO1* present at all five stages. *LFY* transcript levels were more abundant than those of *WAPO1*, and both genes showed lower transcript levels in the apical region at the vegetative stage than at the double-ridge to floret primordia stages (Fig. S3, data S7). These studies indicate that *LFY* and *WAPO1* are present at the same stages of spike development. To refine the localization of *LFY* and *WAPO1* transcripts within the developing spike, we examined their dynamic spatial patterns using smFISH (Fig. 4). For all the smFISH studies, we only compared hybridization signals across developmental stages for individual genes because comparisons of total expression levels among genes are affected by probe sensitivity and can be misleading.

**Fig. 4.**
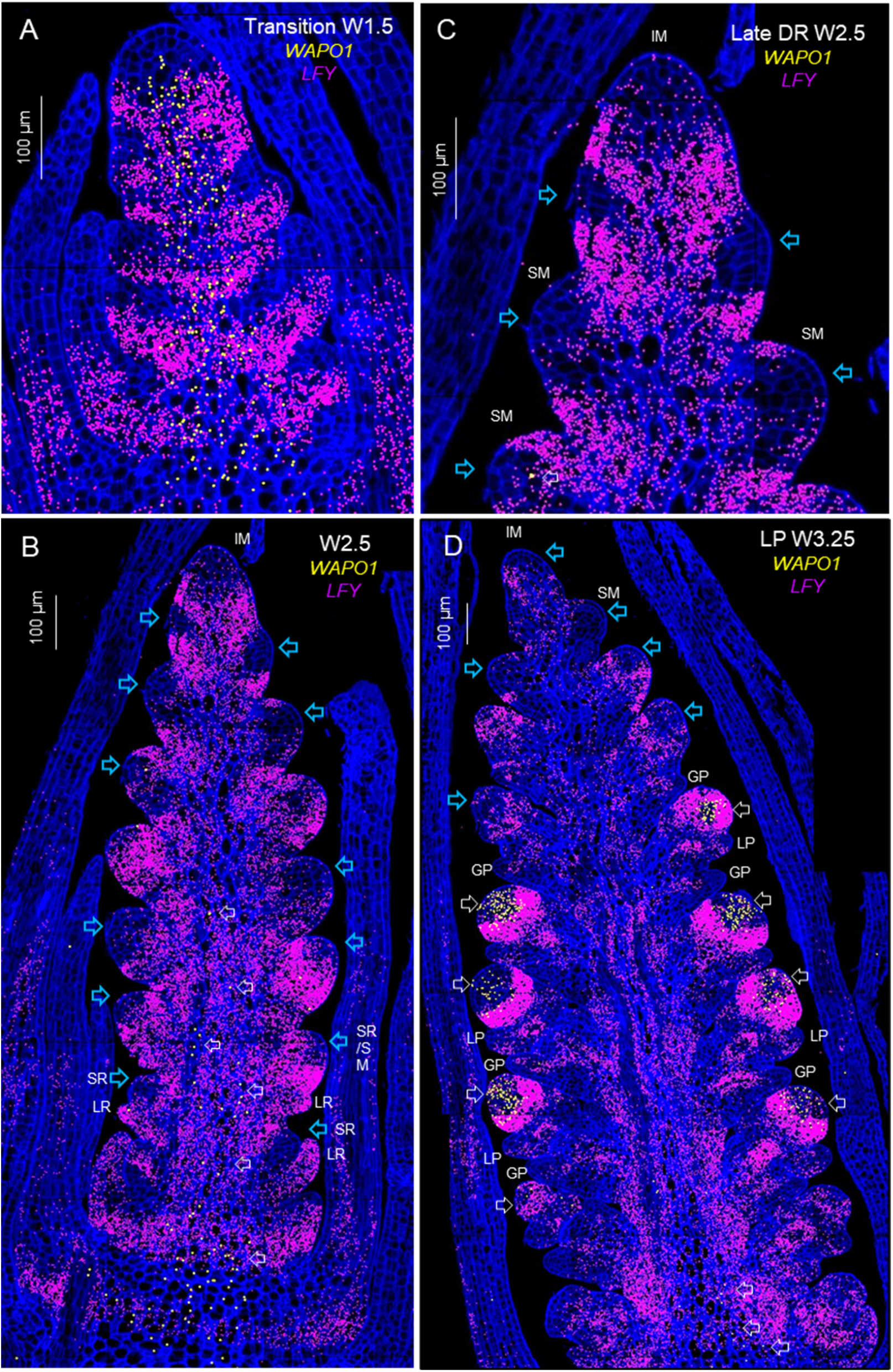
Single-molecule fluorescence in-situ hybridization (smFISH) of *LFY* and *WAPO1* during spike development. Cell walls stained with calcofluor are presented in dark blue. (**A**) Elongated shoot apical meristem transitioning from a vegetative to an inflorescence meristem (IM, W1.5). (**B**) Late double-ridge stage (W2.5). (**C**) Detail of the IM region from (B). (**D**) Lemma primordia stage (W3.25). Blue arrows indicate regions of the SM where *LFY* expression is lower. LR = leaf ridge, SR = spikelet ridge, GP = glume primordium, LP = lemma primordium. Scale bar 100 μm. W = Waddington scale (Waddington *et al*., 1983).

During the transition between the vegetative and reproductive phases (W1.5 in Waddington scale (Waddington *et al*., 1983)), *LFY* transcripts were concentrated in bands radiating from the axis of the elongating shoot apical meristem (SAM) towards the lateral primordia (Fig. 4A). Only few *LFY* transcripts were detected at the tip of the IM at this or later spike developmental stages (Fig. 4C-D). At the late double-ridge stage (W2.5), in the less developed lateral meristems present at the bottom (Fig. 4B) and top (Fig. 4C) of the developing spike, *LFY* expression was stronger at the leaf ridge (also known as lower ridge) than at the spikelet or upper ridge (blue arrows). In the more mature spikelet meristems (SMs) located in the central part of the developing spike, *LFY* expression was abundant in the basal region but low in central-distal regions of the SMs (Fig. 4B, blue arrows), suggesting that low *LFY* levels may favor spikelet development. Some SMs showed a more uniform distribution of *LFY*, but those may be the result of off-centered sections.

We used the gene *FRIZZY PANICLE* (*FZP*, *TraesCS2A02G116900*) as an early marker for the IM→TS transition. At W2.5, *FZP* was not detected in the distal part of the wheat spike and was present only at the axils of the developing glumes in the more mature central spikelets, similar to previously reported results in rice (Komatsu *et al*., 2003). However, at W3.25, when the developing spikes reach the final SNS, *FZP* was detected in the youngest lateral meristems immediately below the IM (Fig. S4, data S8), indicating their transition to glumes and serving as a marker of the IM→TS transition.

At the W3.25 stage, the more developed spikelets at the center of the spike showed glume and lemma primordia (Fig. 4D). In these spikelets, *LFY* transcripts were highly expressed within a narrow band that, in a tridimensional space, may be similar to a bird’s nest located distal to the lemma primordia. This high-expression *LFY* band delimited a distal region of the developing spikelet that has lower *LFY* and higher *WAPO1* hybridization signals (Fig. 4D).

*WAPO1* transcripts were detected at the axis of the developing spike at W1.5, in a region that overlapped with *LFY* (Fig. 4A and S5). However, at later stages (W2.5 and W3.25) *WAPO1* expression was restricted to the base and center of the developing spike, likely in the differentiating vascular tissue (Fig. 4B and D). This distribution is easier to visualize in Fig. S5 that presents *WAPO1* expression alone. At the lemma primordia stage (W3.25), *WAPO1* expression was also detected in the distal part of the more developed spikelets (Fig. 4D and S5C), in agreement with previous *in situ* hybridization results (Kuzay *et al*., 2022). A detail of the *WAPO1* expression domain shows co-localization with *LFY* in multiple cells within the distal region of the developing spikelet, which extends to one - two cell layers into the area of high *LFY* expression (Fig. S6).

In summary, the distinct but partially overlapping expression domains of *LFY* and *WAPO1* in the developing spikelets generate an area of overlap, which likely favors interactions between their encoded proteins and provides important spatial information for normal floral development.

### Spatio-temporal expression profiles of *LFY* and *SQUAMOSA* MADS-box genes

In Arabidopsis, LFY activates the meristem identity genes *AP1* (Parcy *et al*., 1998, Wagner *et al*., 1999) and *CAL* (William *et al*., 2004), so we first tested if the homologous wheat *VRN1* and *FUL2* genes (data S9) were also regulated by *LFY* using qRT-PCR (Fig. S7, data S10). We found no significant differences between *lfy* and the wildtype control for *VRN1* or *FUL2* transcript levels at W2.0, W3.0 or W4.0 (Fig. S7). Analyses of previously published RNAseq data for Kronos spike development (VanGessel *et al*., 2022) showed that *VRN1* is induced earlier and is expressed at higher levels than the other two *SQUAMOSA* genes, with *FUL2* expressed at higher levels than *FUL3* (Fig. S8A). In the same RNAseq study, *LFY* was expressed at low levels in the vegetative meristem (W1.0) and increased rapidly during W2.0 and W3.0 (Figs. S3B and S8A).

We then compared the smFISH spatial and temporal expression profiles of *VRN1* and *FUL2* during spike development. At the late vegetative stage (W1.0), the hybridization signal of *VRN1* was relatively low and *FUL2* was not detected in the apical meristem (Fig. 5A). The signal for both genes increased during the early transition to the reproductive stage, although *FUL2* remained low (Fig. 5B, W1.5). These results were consistent with the RNAseq data (Fig. S8A). At later stages (W2.5 and W3.25), *VRN1* and *FUL2* were both highly expressed in the IM and young lateral SMs (Fig. 5C-D). In the more developed spikelets, located at the center of the developing spike, *VRN1* and *FUL2* expression was stronger at the glume and lemma primordia than in the distal region (Fig. 5D, W3.25), which overlapped with the *WAPO1* expression domain (Fig. 4D). *FUL3* showed a similar spatial expression profile as *FUL2* and it is presented separately (Fig. S9) because of its lower expression levels in the RNAseq data (Fig. S8A) and limited impact on SNS (Li *et al*., 2019).

**Fig. 5.**
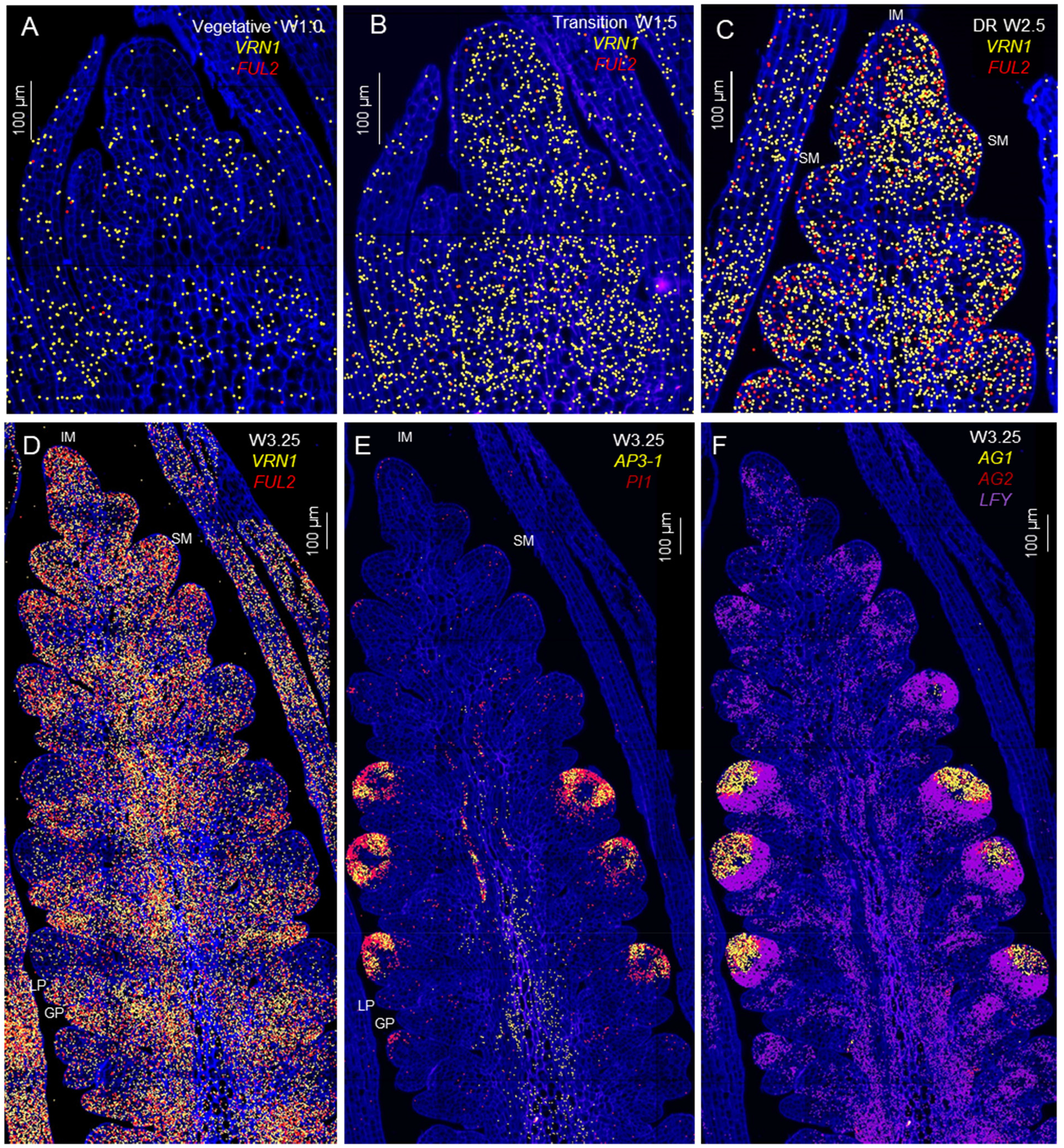
Transcription profiles of MADS-box floral genes during wheat spike development. (**A-B**) Relative distribution of *VRN1* and *FUL2*. (**A**) Late vegetative SAM. (**B**) Transitioning SAM. (**C**) *VRN1* and *FUL2* at late double-ridge. (D-F) Lemma primordia stage. (**D**) *VRN1* and *FUL2*. Class-B MADS-box genes (**E**) *AP3-1* and PI1 and (**F**) *AG1* and *AG2*. IM=inflorescence meristem, SM=spikelet meristem, GP=glume primordium, LP=Lemma primordium. Gene identifications and rice orthologs are in data S9.

To quantify *VRN1*, *FUL2* and *LFY* expression changes in the distal part of the developing spike between the late double ridge stage (W2.5) and the start of the transition to a terminal spikelet (W3.25), we calculated their hybridization signal per 100 μm^2^ (signal density). In both the IM and the IM plus the two youngest lateral meristems (IM+2LM), we observed a 3- to 4-fold increase in *FUL2* signal density (Fig. S8B) and a 53-69% increase in *VRN1* between W2.5 and W3.25, though only the differences for *FUL2* were significant (Fig. S8C, data S8). Similar results were obtained when the *VRN1* and *FUL2* hybridization signals were normalized using the *CDC20* signal (data S8). In the same tissue sections, we detected a 78% decrease in *LFY* signal density (Fig. S8D), and these changes were significant or highly significant depending on the normalization method used (data S8).

Analyses of the ratios between the *SQUAMOSA* and *LFY* signals showed that the *FUL2/LFY* ratio increased more than 20-fold between W2.5 and W3.25 in both the IM and IM+2LM (Fig. S8E, data S8). Similarly, *VRN1/LFY* ratios increased 8-fold between the same developmental stages (Fig. S8E, data S8). The *SQUAMOSA/LFY* ratios are independent of the normalization method used, and since they are determined in the same tissue sections, they provide the best evidence of a significant increase in the expression of the *SQUAMOSA* genes relative to *LFY* in the distal part of the developing spike at the time of the IM→TS transition.

### Spatial expression profiles of floral organ identity genes

We also characterized the spatial distribution of MADS-box genes involved in floral organ development (data S9). The hybridization signals of class-B (*AP3-1* and *PI1*, Fig. 5E), class-C (*AG1*and *AG2*, Fig. 5F) and class-E (*SEP3-1* and *SEP3-2*, Fig. S10) floral organ identity genes were concentrated in a distal region of the developing spikelets that mostly overlapped with the expression of *WAPO1* (Fig. 4D and 5D). *SEP1-2*, *SEP1-4*, and *SEP1-6* were expressed outside of the region where the two *SEP3* were expressed (Fig. S10), suggesting functional divergence between the *SEP1* and *SEP3* genes in wheat.

Finally, we used qRT-PCR to characterize the effect of the *lfy* mutation on the expression of the floral organ identity genes in the wheat developing spike at W4.0, when these genes are highly expressed (Kuzay *et al*., 2022). The *lfy* mutant showed a significant downregulation of *AP3-1* and *PI1* (Fig. S11A), *AG1* (Fig. S11B), *SEP3-1* and *SEP3-2* (Fig. S11C) relative to the wildtype (data S11). Taken together these results indicate that *LFY* plays an important role in the direct or indirect regulation of the floral organ identity genes.

### Genetic interactions between *LFY* and class-A MADS-box genes

Given the opposite effects of *LFY* and the MADS-box genes *VRN1* and *FUL2* on SNS and their opposite expression changes during the IM→TS transition, we examined their genetic interactions for this trait. We crossed a plant homozygous for *lfy-A* and heterozygous for *lfy-B* with mutants homozygous for *vrn1* and *ful2*-A but heterozygous for *ful2*-B and, in the progeny, selected sister plants homozygous for the four gene combinations for each gene (WT, *lfy*, *vrn1*, *lfy vrn1* and WT, *lfy*, *ful2*, *lfy ful2*).

A factorial ANOVA including the four homozygous *VRN1-LFY* combinations showed highly significant effects on SNS for both *VRN1* and *LFY*, and a highly significant interaction between these two genes (*P*<0.0001, Fig. 6A, data S12). The effect of *LFY* on SNS was stronger in the *vrn1* mutant (10.3 spikelets) than in the presence of the functional *Vrn1* allele (5.2 spikelets). In contrast, the effect of *VRN1* on SNS was stronger in the presence of the functional *LFY* allele (9.1 spikelets) than in the presence of the *lfy* combined mutant (4.0 spikelets, Fig. 6A). *VRN1* also showed highly significant effects on the number of leaves and heading time, similar to previous studies (Li *et al*., 2019), while *LFY* showed no-significant differences for these traits in the presence of the *Vrn1* or *vrn1* alleles. No significant interactions between these two genes were detected for these two traits (Fig. 6B-C, data S12).

**Fig. 6.**
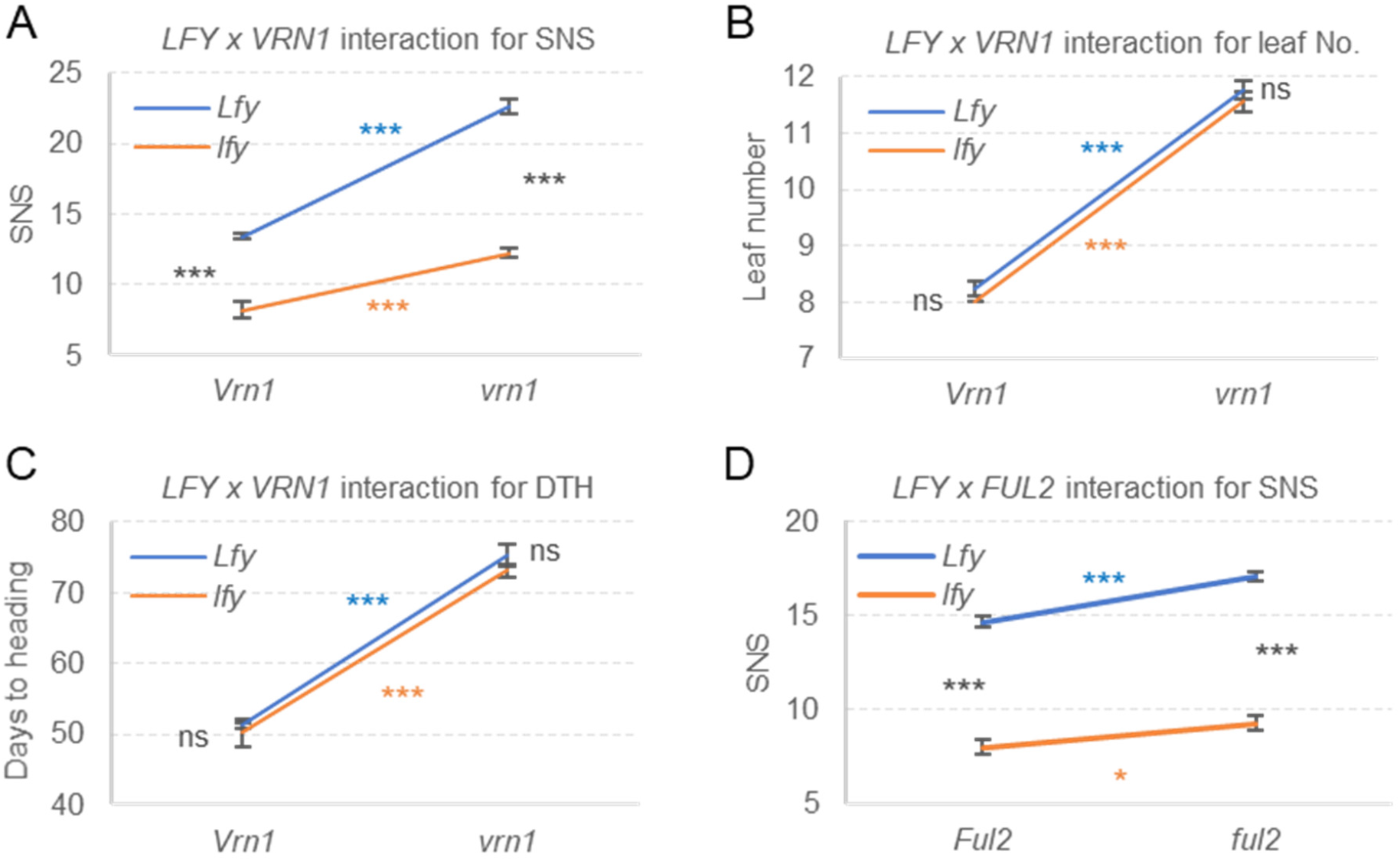
**Genetic interactions between *LFY* and *SQUAMOSA* MADS-box genes *VRN1* and *FUL2***. (**A-C**) Interactions graphs between *LFY* and *VRN1* for (**A**) SNS (total N=34), (**B**) leaf number (total N=34), and (**C**) days to heading (total N=34). (**D**) Interaction between *LFY* and *FUL2* for SNS (total N=54). In the interaction graphs, parallel lines indicate additive effects and non-parallel lines reflect interactions. *P* values of simple effect (data S12). ns=not significant, *=*P<*0.05, **=*P<*0.01, and ***=*P<*0.001. Error bars are SEM. Raw data in data S12.

The effect of *FUL2* on SNS was smaller than the effect of *VRN1* (Fig. 6D, adjusted means from two experiments) but the main effects of both *LFY* and *FUL2* were still highly significant in both experiments. The contrasts for the four single effects were also consistent between the two *FUL2* experiments (data S12) and also with the interactions between *LFY* and *VRN1*: a stronger effect of *LFY* on SNS in the presence of the mutant *ful2* allele than in the presence of the wildtype *Ful2* allele, and a stronger effect of *FUL2* in the presence of wildtype *LFY* allele than in the presence of *lfy* (data S12). The *FUL2* x *LFY* interaction was significant only in the second experiment with a higher number of replications (*P*=0.0175). In summary, these interactions suggest the existence of a cross-talk between the *SQUAMOSA* and *LFY* genes in the regulation of SNS.

## DISCUSSION

### Similarities and differences in *LFY* function between Arabidopsis and wheat

The most conserved functions of *LFY* across the flowering plants are those associated with the regulation of organ identity in the three inner floral whorls, which include pistils, stamens and petals in eudicot or lodicules in grass species (Yoshida, 2012). *LFY* mutations have more limited effects on bracts (lemmas in grasses) or on the outermost floral whorls, including sepals in Arabidopsis and paleas in grasses. However, the first floret of the basal spikelet frequently showed a bifurcated palea (Fig. 1F.), suggesting that interactions between *LFY* and genes expressed in the base of the wheat spike regulate palea development.

In the grass species, similar defects in the inner floral organs have been observed for wheat *lfy* mutants (Fig. 1F and data S2, this study), rice *apo2* mutants (Ikeda-Kawakatsu *et al*., 2012), barley *multiovary 5* mutants (Selva *et al*., 2021), and maize *zfl1 zfl2* mutants (Bomblies *et al*., 2003). These defects include fused organs, reduced number and altered morphology of lodicules (including transformation into bracts and fusions with stamens), reduced number of stamens and increased number of pistils. Fused lodicules with stamens and homeotic conversions of stamens into pistils were observed frequently in the wheat *lfy* mutant (Fig. 1F). The Arabidopsis strong *lfy* mutants fail to develop flowers, but weak *lfy* mutants show petals transformed into small sepals or mosaic organs, reduced stamen numbers and increased pistil numbers (Huala and Sussex, 1992, Weigel *et al*., 1992), similar to the grass species.

Despite the conserved roles of *LFY* in floral organ development, there are also important differences in *LFY* functions between Arabidopsis and grasses. First, strong *lfy* mutations in Arabidopsis result in the replacement of most flowers by shoots subtended by cauline leaves and the few observed late flowers exhibit intermediate inflorescence characteristics (Schultz and Haughn, 1991, Weigel *et al*., 1992). In contrast, floret meristems initiate normally in *lfy* in wheat (Fig. 1F) and other grasses (Bomblies *et al*., 2003, Ikeda-Kawakatsu *et al*., 2012), and defects appear only later at the inner floral whorls. These results indicate that *LFY* is required to confer the initial floral meristem identity in Arabidopsis but not in the grass species.

This difference likely contributes to the opposite functions of *LFY* in inflorescence development in these species. In Arabidopsis, constitutive expression of *LFY* (*35S:LFY*) results in conversion of both apical and axillary meristems into terminal flowers, demonstrating that *LFY* is a limiting factor defining when and where flowers are produced (Weigel and Nilsson, 1995). In contrast, constitutive expression of *LFY* in wheat increases the number of lateral spikelets (Fig. 2).

Mutations in *LFY* also result in contrasting effects in eudicots and grasses inflorescence development. Weak *lfy* mutants delay the formation of flowers and increase the number of secondary branches in Arabidopsis and other eudicot species (Coen *et al*., 1990, Schultz and Haughn, 1991, Weigel *et al*., 1992, Souer *et al*., 1998, Molinero-Rosales *et al*., 1999). In contrast, *LFY* loss-of-function mutations result in significant reductions in SNS in wheat (Fig. 1B-E), and in the number of branches in rice panicles (Ikeda-Kawakatsu *et al*., 2012) and maize male inflorescences (Bomblies *et al*., 2003).

In summary, *LFY* has a similar role on floral organ development in Arabidopsis and grasses, but plays different roles in floral meristem identity and inflorescence development.

### LFY and WAPO1 jointly regulate floret and spike development

LFY and WAPO1 proteins physically interact with each other in wheat (Fig. 3A), barley (Selva *et al*., 2021), rice (Ikeda-Kawakatsu *et al*., 2012) and Arabidopsis (Chae *et al*., 2008). In Arabidopsis, the LFY-UFO complex binds to a different set of genes than LFY alone (Rieu *et al*., 2023b), including the class-B genes *AP3-1* and *PI.* In addition to the class-B genes, *ufo* mutations in other eudicot species show altered expression of class-C genes (snapdragon), class- E genes (cucumber) and both class-C and -E genes (petunia, reviewed in (Rieu *et al*., 2023a)).

The simultaneous regulation of class-B, -C and -E MADS-box genes seems to be conserved in the temperate grasses, where *lfy* mutations in wheat (Fig. S11) and barley (Selva *et al*., 2021), and *wapo1* mutants in wheat (Kuzay *et al*., 2022) have all been associated with the downregulation of class-B (*AP3-1* and *PI1*), class-C (*AG1*) and class-E (*SEP3*) floral organ identity genes. This result explains the similar floral defects observed in the wheat *lfy* (Fig. 1F) and *wapo1* mutants (Kuzay *et al*., 2022), and indicates that both LFY and WAPO1 are required for the proper regulation of these floral organ identity genes.

In addition to their roles in floral development *LFY* and *WAPO1* jointly regulate SNS. The *wapo1*, *lfy* and combined *lfy wapo1* mutants all show similar reductions in SNS (Fig. 3B). Similar reductions in the number of panicle branches are also observed in the *apo1*, *apo2* and combined *apo1 apo2* mutants in rice (Ikeda-Kawakatsu *et al*., 2012). In this study, we show that the reduction in wheat SNS in the *lfy* and *wapo1* mutants is the result of a reduction in the rate of SM formation per day rather than a change in the timing of the IM→TS transition. Interestingly, in tomato overexpression of either *LFY* (= *FALSIFLORA*) or *UFO* (= *ANANTHA*) can transform multiflowered inflorescences into solitary flowers (MacAlister *et al*., 2012). By contrast, heterochronic shifts during meristem maturation of *UFO* and few other flowering regulators can modulate inflorescence complexity (Lemmon *et al*., 2016). These results suggest that LFY and UFO homologs can affect meristem maturation rates in both tomato and wheat.

In summary, *LFY* and *WAPO1* act jointly to regulate both inflorescence architecture and floral organ development in wheat, and likely in other plant species.

### Interactions between *LFY* and *SQUAMOSA* genes modulate their opposite effects on the IM development

MADS-box transcription factors act as master regulators of developmental switches and organ specification, with meristem identity genes from the *SQUAMOSA* clade playing essential roles in the initiation of flower development. Among them, AP1 can modulate chromatin accessibility and facilitate access of other transcriptional regulators to their target gene, suggesting that it acts as a pioneer transcription factor (Pajoro *et al*., 2014). In the wheat *vrn1 ful2 ful3* combined mutant, lateral spikelets are completely transformed into vegetative tillers subtended by leaves (Li *et al*., 2019), whereas the flowers in the Arabidopsis *ap1 cal ful* triple mutant are transformed into leafy shoots (Ferrándiz *et al*., 2000). LFY is also a pioneer transcription factor that can bind nucleosomes in closed chromatin, displace H1 linker histones and recruit the SWI/SNF chromatin-remodeling complex, permitting the binding of other transcription factors (Jin *et al*., 2021, Lai *et al*., 2021, Yamaguchi, 2021).

In Arabidopsis, LFY positively regulates the expression of *AP1* and *CAL* (Parcy *et al*., 1998, Wagner *et al*., 1999, William *et al*., 2004), thereby indirectly regulating their downstream targets. *LFY* also directly regulates hundreds of genes independently of *AP1* and *CAL* (Goslin *et al*., 2017). However, induction of *LFY* in *ap1 cal* mutants is insufficient to rescue the limited and late formation of flowers observed in this mutant and, instead, results in a lower proportion of plants with flowers than in the control (Goslin *et al*., 2017). This last result indicates that, in the absence of *AP1* and *CAL, LFY* can inhibit flower formation in Arabidopsis, similar to its function in the grass species. These results also suggest that the ability of LFY to directly regulate the *SQUAMOSA* genes in Arabidopsis but not in wheat (Fig. S7) likely contributes to the opposite functions of *LFY* in the regulation of IM development between these species.

Approximately 200 genes have been identified in Arabidopsis as high-confidence direct targets of both LFY and AP1 (Winter *et al*., 2015). Many of the shared genes directly regulated by the induction of *AP1* or *LFY* show changes in expression with identical directionality, but some of them are regulated in opposite directions, including several key regulators of floral initiation (Goslin *et al*., 2017). These common gene targets can contribute to the epistatic genetic interactions between *LFY* and *AP1* and to their ability to coordinate the transcriptional programs required for flower initiation and early flower development in Arabidopsis.

Genes that are directly regulated by both LFY and SQUAMOSA are likely to also exist in wheat, and may contribute to the significant genetic interaction for SNS observed in this study between *LFY* and both *VRN1* (Fig. 6A) and *FUL2* (Fig. 6D). The net effect of the *LFY x VRN1* interaction, 5.2 spikelets per spike, was smaller than the maximum differences of 14.4 spikelets observed between the plants carrying the *vrn1 Lfy* combination (22.6 spikelets/spike) and those carrying the *Vrn1 lfy* combination (8.2 spikelets/spike, data S12). These results indicate that the interaction between *LFY* and *SQUAMOSA* explains only part of the variation in SNS.

In summary, *lfy* mutations have opposite effects on SNS than *vrn1* or *ful2* mutations, and those effects are mostly additive. However, significant genetic interactions between *LFY* and the *SQUAMOSA* genes contribute to the observed differences in SNS in the different mutant combinations.

### *LFY* and *WAPO1* show dynamic spatio-temporal expression patterns during wheat spike and spikelet development

#### Spike development

From the beginning of the wheat spike development, *LFY* expression is not uniform, with higher expression levels at the lower or leaf ridge than at the upper or spikelet ridge (Fig. 4A-C), a pattern reported previously in wheat by *in situ* hybridization (Shitsukawa *et al*., 2006). This spatial differentiation continues in the early spike development (W2.5), where *LFY* is abundant in the proximal region of the young SMs but rare in their distal region (Fig. 4B-C). A similar pattern was also reported in the SMs of young barley spikes (Zhong *et al*., 2021) and in the primary and secondary branch meristems in the developing rice panicles (Kyozuka *et al*., 1998, Miao *et al*., 2022). These results suggest a conserved *LFY* spatial pattern in developing grass inflorescences and highlight the importance of reduced *LFY* levels in initiating spikelet development.

In wheat, the SM regions with low *LFY* expression (Fig. 4B-C) show high levels of *VRN1* and *FUL2* transcripts (Fig. 5C-D) and, therefore, high *SQUAMOSA / LFY* ratios. We hypothesize that this change in the balance between SM promoting and repressing pioneer transcription factors is critical for marking the regions where the lateral spikelets will develop. This hypothesis is also supported by the drastic changes in the relative smFISH signal densities of these genes in the IM and two youngest lateral meristems between the early stages of spike development (W2.5) and the time of the IM→TS transition (determined by the presence of *FZP*, W3.25 Fig. S4). In the IM, the *VRN1/LFY* ratio increased more than 8-fold and the *FUL2/LFY* ratio increased more than 25-fold between W2.5 and W3.25 at the initiation of the terminal spikelet (Fig. S8 and data S8).

Changes in gene dosage for *LFY* (Fig. S1) and the *SQUAMOSA* genes result in changes in SNS, confirming the importance of their relative transcript levels on spike development. The single *lfy- A* and *lfy-B* mutants show intermediate reductions in SNS relative to the combined *lfy* mutant (Fig. S1), whereas mutations in *FUL2* result in smaller increases in SNS than mutations in *VRN1*, which is expressed at higher levels than *FUL2* in the IM (Fig. S8A and data S8) (Li *et al*., 2019). Interestingly, four recently cloned genes affecting SNS in wheat including *WAPO1* (Kuzay *et al*., 2022), *FT-A2* (Shaw *et al*., 2019, Glenn *et al*., 2022), *bZIPC1* (Glenn *et al*., 2023) and *SPL17* (Liu *et al*., 2023) show potential connections with the regulation of *VRN1* or *LFY* (Fig. S12). In summary, we propose that the modulation of *VRN1* or *LFY* transcript levels in the IM plays an important role in the determination of SNS in wheat.

*LFY* was co-expressed with *WAPO1* in the IM at the early stages of spike development (Fig. 4A) but not at later stages (Fig. 4B-C and Fig. S5B-C). This early co-localization in the IM seems to be sufficient to explain the similar reduction in SNS observed in the *lfy* and *wapo1* mutants. Both mutants were associated with similar reductions in the rate of SM formation relative to the wildtype, rather than by a change in the timing of the IM→TS transition (Fig. 3C-D). Since these differences were evident from the earliest stages of spikelet development, when both *LFY* and *WAPO1* were co-expressed in the IM (Fig. 4A), we hypothesize that the transient formation of the LFY-WAPO1 complex is sufficient to activate gene expression networks that accelerate the rate of SM initiation.

#### Spikelet development

The smFISH studies also provided insights into the dynamic spatial distribution of *LFY*, *WAPO1*, and the floral organ identity genes during spikelet development. In the more developed central spikelets at W3.25, *LFY* was highly expressed in a narrow nest-shaped region distal to the lemma primordia, delimiting a distal spikelet meristem region with lower *LFY* expression (Fig. 4D).

This intense *LFY* expression band has been also observed distal to the lemma primordia of the second and third florets by *in situ* hybridization in more developed wheat spikelets (Shitsukawa *et al*., 2006), highlighting its importance for normal floret development.

Within the distal region of the developing spikelets, *WAPO1* was co-expressed with *LFY* in multiple cells including one or two cell layers within the region of high *LFY* expression (Fig. 4D and S6). Within this overlapping region LFY and WAPO1 proteins may have a higher chance to interact with each other, providing valuable spatial information to the floral organ identity genes. This hypothesis is indirectly supported by similar reductions in the expression levels of the floral organ identity genes and similar floral abnormalities in both the *wapo1* and *lfy* mutants (Fig. 1F, S11, (Kuzay *et al*., 2022)).

In Arabidopsis, the UFO-LFY complex regulates a different set of gene targets than LFY alone (Rieu *et al*., 2023b), and both genes are required for the correct regulation of the floral organ identity genes *AP3* (Chae *et al*., 2008, Rieu *et al*., 2023b) and *PI* (Honma and Goto, 2000). These results are consistent with the down-regulation of the wheat floral organ identity genes in the *lfy* and *wapo1* mutants, and with the overlap between the expression domains of the wheat floral organ identity genes (Fig. 5 and S10) and *LFY* - *WAPO1* co-expression in the distal part of the wheat developing spikelets.

In summary, this study shows that *LFY* plays important roles in both floral organ and spike development. The dynamic spatial and temporal expression profiles of *LFY* and *WAPO1* in the developing spikelets correlate well with the function of these genes in the regulation of floral organ identity genes and normal floret development. In addition, these genes regulate the rate of SM formation, and interact with the *SQUAMOSA* genes in the regulation of SNS. Therefore, natural or induced variation in these genes can be used to improve this important agronomic trait in a crop that is central for global food security.

## MATERIALS AND METHODS

### Ethyl methanesulfonate (EMS) induced *LFY* mutants and their interactions with *VRN1*, *FUL2* and *WAPO1*

We screened the sequenced tetraploid wheat variety Kronos population (Krasileva *et al*., 2017) by BLASTN to identify loss-of-function mutations in the *LFY-A1* and *LFY-B1* homeologs. To reduce background mutations, the *lfy-B* mutant was backcrossed twice to Kronos and then to the *lfy-A* mutant. The double mutant was backcrossed once to Kronos and, among the progeny, homozygous sister lines were selected for the four possible homozygous combinations, including the *lfy-A lfy-B* combined mutant (henceforth, *lfy*). This line is BC1 for *lfy-A* and BC2 for *lfy-B* so it is referred to as BC1-2. Genome specific markers for the *lfy-A* and *lfy-B* mutations were designed and used to genotype plants during backcrossing and combination with other mutations described below. Primers are listed in data S1.

To study the interactions between *LFY* and other spike development genes, we intercrossed the *lfy* combined mutant with previously developed Kronos lines homozygous for loss-of-function EMS or CRISPR mutations in both genomes of *VRN1* (Chen and Dubcovsky, 2012), *FUL2* (Li *et al*., 2019), or *WAPO1* (Kuzay *et al*., 2022). These lines had at least two backcrosses to the parental line Kronos to reduce the number of background mutations. For these crosses, we used a line heterozygous for one of *LFY* mutation to restore fertility, and molecular markers to select the four possible homozygous combinations in the F2 progeny of each cross.

### Plant growth and phenotypic characterization

Plants were stratified for 2-days at 4 °C in the dark and then planted in growth chambers (PGR15, Conviron, Manitoba, Canada). Lights were set to 350 μmol m^-2^ s^-1^ at canopy level. Plants were grown under inductive 16-h long days with temperatures set during the day to 22°C and during the night to 17°C. Relative humidity in growth chambers was maintained at 60-70% throughout the duration of the experiments. Heading time was recorded as the number of days from germination to full emergence of the spike from the leaf sheath. Spikelet number per spike (SNS) was measured at maturity from the main tiller.

### Generation of the wheat transgenic lines overexpressing *LFY*

We cloned the *LFY-A* coding regions by PCR from cDNA derived from Kronos developing spikes using primer LFY-A-GW-F combined with LFY-A-GW-R2 (listed in data S1). We then recombined it into pDONR/zeo entry vector using Life Technologies BP Clonase II following the manufacturer’s protocol. The pDONR/zeo vector containing the *LFY-A* coding region was next recombined into the Japan Tobacco pLC41 vector downstream of the maize *UBIQUITIN* promoter using Life Technologies LR Clonase II to generate a construct expressing *LFY* with a C-terminal 3xHA tag, which was verified by Sanger sequencing at each cloning step. The T-DNA binary construct was transformed into the wheat variety Kronos using *Agrobacterium-* mediated transformation (EHA105) at the UC Davis Plant Transformation Facility as described before (Debernardi *et al*., 2020b). The presence of the *LFY* transgene was determined with primers LFY-Genotyping-R5 and UBI-F2 listed in data S1.

*LFY* transcript levels were determined by qRT-PCR using primers LFY_qPCR_F and LFY_qPCR_R and *ACTIN* as endogenous control (data S1). For qRT-PCR experiments, RNA was extracted using the Spectrum Plant Total RNA Kit (Sigma-Aldrich, St. Louis, MO, USA) or as previously described by (Ream *et al*., 2014). One μg of RNA was treated with RQ1 RNase-Free DNase (Promega, M6101) first and then used for cDNA synthesis with the High-Capacity cDNA Reverse Transcription Kit (Applied Biosystems, Foster City, CA, USA). The qRT-PCR experiments were performed using Quantinova SYBR Green PCR kit (Qiagen, 208052) in a 7500 Fast Real-Time PCR system (Applied Biosystems). Transcript levels for all genes are expressed as linearized fold-*ACTIN* levels calculated by the formula 2^(*ACTIN*^ ^CT –^ *^TARGET^* ^CT)^ ± standard error (SE) of the mean.

### Single-molecule fluorescence *in-situ* hybridization (smFISH)

We used the Molecular Cartography^TM^ technology from Resolve BioSciences combinatorial smFISH. Wheat shoot apical meristems were collected from the vegetative to the spike lemma primordia stage. The samples were immediately fixed in 4% PFA after harvest, dehydrated, and embedded in paraffin. Sections from the central plane of the developing spikes (10µm-thick) were placed on the slides and dried overnight at 37°C, followed by a 10-minute bake at 50°C. The sections were then deparaffinized, permeabilized, and refixed according to the user guide. After complete dehydration, the sections were mounted using SlowFade-Gold Antifade reagent, covered with a thin glass coverslip, and sent to Resolve BioSciences on dry ice for analysis as described in our previous study (Glenn *et al*., 2023).

Probes were designed using Resolve BioSciences’ proprietary design algorithm and gene annotations from Chinese Spring RefSeqv1.1. To identify potential off-target sites, searches were confined to the coding regions. Each target sequence underwent a single scan for all k-mers, favoring regions with rare k-mers as seeds for full probe design. For each of the wheat genes selected for smFISH probe design (data S9), we selected the homoeolog expressed at higher levels in a Kronos transcriptome including different stages of spike development (VanGessel *et al*., 2022), and provided Resolve BioSciences their respective homeologs to be excluded in their quality control for primer specificity performed against all the coding sequences of the wheat genome (Ref Seq v1.1). Therefore, probes are not genome-specific and may detect both homeologs for each gene (Catalogue No. PGGS, all these probes are part of kit number K7128).

Imaging and image processing was performed as described before (Glenn *et al*., 2023). Final image analysis and quantification of dot density was performed in ImageJ using the Polylux tool plugin to examine specific Molecular Cartography^TM^ signals.

### Co-immunoprecipitation (Co-IP) assay and western blotting

To test the physical interaction between WAPO and LFY, we performed Co-IP experiments in wheat leaf protoplasts using a method described previously (Zhang *et al*., 2023), with minor modifications. The *WAPO1* coding region was initially synthesized by Genewiz into the pUC57 vector, amplified with primers WAPO1_BP_F and WAPO1_BP_R (data S1), cloned into pDONR/zeo vector using Life Technologies BP Clonase II, and recombined into the Japan Tobacco pLC41 vector downstream of the maize *UBIQUITIN* promoter with a C-terminal 4xMYC tag (for transgenic experiments). Next, we switched both *UBI:WAPO1-MYC* and *UBI:LFY-HA* from the pLC41 binary vector to the smaller pUC19 vector to enhance the transfection efficiency of the protoplasts. The *UBI:WAPO-MYC* and *UBI-LFY-HA* DNA fragments were cleaved using restriction enzymes *Hind*III and *Spe*I, gel purified, and then ligated with the *Hind*III-*Xba*I linearized pUC19 vector (*Spe*I and *Xba*I create compatible ends). Both constructs were verified by restrictions and Sanger sequencing.

We transformed Kronos leaf protoplasts with 50 μg of each of the *UBI:LFY-HA* and *UBI: WAPO1-MYC* plasmids in 50 ml tubes containing 2 ml of protoplast (roughly 0.5 x 10^6^ cell per mL). As negative controls, we performed separate transformations including only one of the two plasmids. After transformation, protoplasts were resuspended in 5 ml W5 buffer and incubated in a 6-well plate at room temperature overnight. Total protein was extracted with 1 ml of IP lysis buffer (25 mM Tris-HCl pH7.5, 150 mM NaCl, 1 mM EDTA, 1% NP-40 substitute, 5% glycerol and 1 x Protease Inhibitors). Part of the protein extract was set aside as input control (50 μl), and the rest was used for co-immunoprecipitation using Pierce Anti-HA Magnetic Beads (ThermoFisher Cat. 88836) by gentle agitation on a tube rotator for 30 min at room temperature with additional 1x proteinase inhibitors. Proteins were eluted by boiling the beads in 50 μl 1x Laemmli sample buffer for 10 min.

For Western Blot, half of the Co-IP elution and 50 μg of input for each sample were loaded onto a 12% SDS-PAGE gel. After protein was transferred to a PVDF membrane using the Bio-Rad Trans-Blot Turbo Transfer System (Cat. 1704150), the membrane was blocked with 1x TBST buffer containing 5% non-fat milk for 1 h at room temperature. Anti-cMyc-peroxidase monoclonal antibody (Roche 11814150001) was added at a dilution of 1:10,000 and was incubated at room temperature for 1h. After four 10 min washes using 1x TBST buffer, the signals were developed using SuperSignal West Femto Maximum Sensitivity Substrate (ThermoFisher, Cat. 34096). After imaging, the membrane was stripped with mild stripping buffer (1.5% glycine, 0.1% SDS, 1% Tween 20, pH 2.2), re-blocked for 1 h at room temperature, and then probed with anti-HA-peroxidase at a dilution of 1:2500 (Roche 12013819001) for 1 h at room temperature.

## ACKNOWLEDGEMENTS

The authors thank Stephen Pearce (Rothamsted Research) and Youngjun Mo (JeonBuk National University) for identifying the *LFY-B* mutant, developing markers and the initial backcrossing to Kronos. We thank Maria von Korff (Heinrich Heine University Düsseldorf) for a valuable revision of the manuscript and for her supervision of Tianyu Lan. We also thank Dr. Kun Li (UC Davis) for her help with multiple experiments, Dr. Junli Zhang (UC Davis) for valuable suggestions, and the UC Davis transformation facility for the generation of the transgenic plants.

## AUTHOR CONTRIBUTIONS

Conceptualization: JD Methodology: HL, FP, CL, AJ Investigation: FP, HL, CL, TL, CT, JMD, AJ Visualization: FP, HL,CL Supervision: JD, DPW, JMD, CL Writing—original draft: FP Writing—review & editing: FP, HL, CL, DPW, JMD, AJ, JD

## DATA AVAILABILITY

All data used in this study are available in the supplementary materials. Kronos mutants K2613 for *lfy-A* and K350 for *lfy-B* are available from the authors upon request without any restrictions for use or from the Germplasm Resources Unit (GRU) at the John Innes Centre. Images for the spike sections used in the smFISH and hybridization coordinates for the genes presented in this study are available at https://dubcovskylab.ucdavis.edu/content/spatial-transcriptomic.

## Supplementary Materials

### Supplementary Figures S1 to S12 are included in a separate PDF

Supplementary data S1 to S12 (listed below) are included in Excel file Supplementary_Data.xlsx **data S1**. Primers used in this study. **data S2**. Supporting data for Fig. 1C-F. **data S3**. Supporting data for Fig. S1. **data S4**. Supporting data for Fig. S2. **data S5**. Supporting data for Fig. 2. **data S6**. Supporting data for Fig. 3B. **data S7**. Supporting data for Fig. S3. **data S8**. Supporting data for Fig. S8. **data S9**. Genes used in the smFISH experiments. **data S10**. Supporting data for Fig. S7. **data S11**. Supporting data for Fig. S11. **data S12**. Supporting data for Fig. 6.

**Fig. S1.**
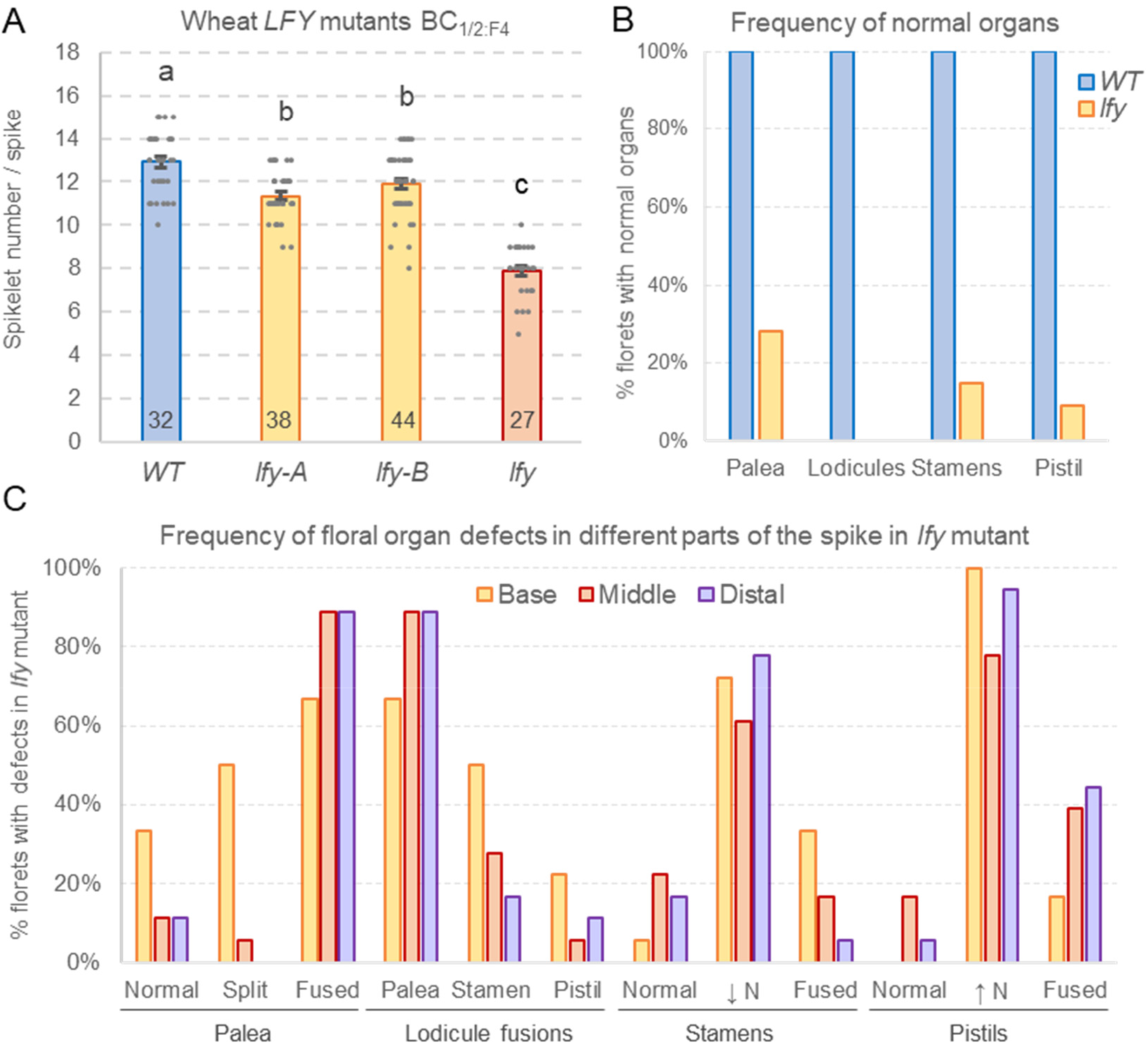
**Effect of *LFY* mutants on spikelet number per spike (SNS) and floral defects**. (**A**) Effect of single (*lfy-A* and *lfy-B*) and combined *lfy* mutants on SNS. Numbers inside the bars indicate number of individual plants measured. Different letters on top of the bars indicate significant differences based on two-tail Tukey test (*P <* 0.05). Raw data and statistics are available in data S3. (**B**) Frequency of florets with normal and abnormal floral organs in Kronos wildtype and *lfy* mutant ( n = 54 florets). Abnormalities include increased or reduced number, morphological changes and fusions with other organs. (**C**) Frequency of abnormalities in the different floral organs in *lfy* mutant, based on 18 florets from the base of the spike, 18 from the middle and 18 from the distal part of the spike. Half of the florets correspond to the 1^st^ floret and the other half to the 2^nd^ floret in the selected spikelets. Raw data are available in data S2.

**Fig. S2.**
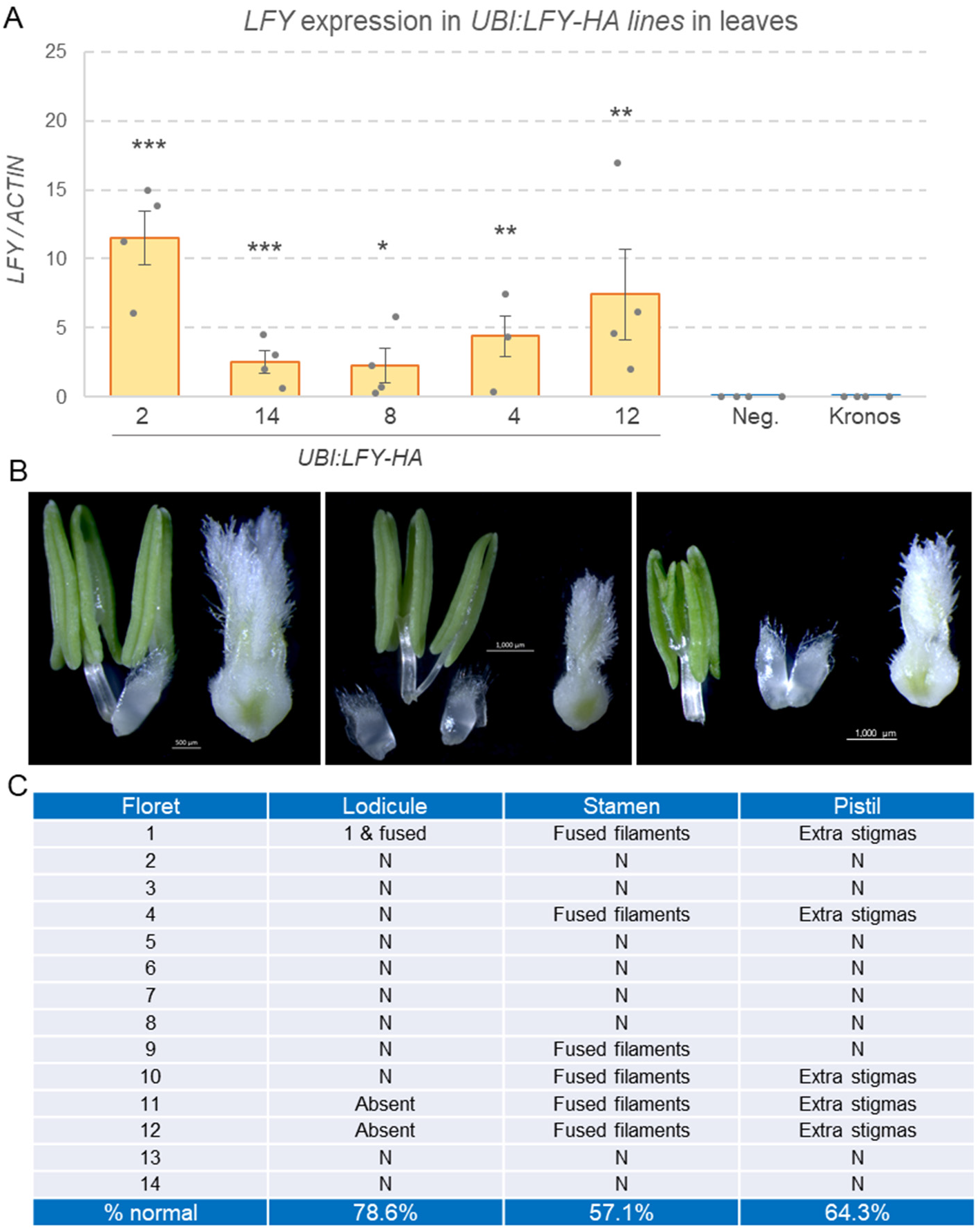
Transgenic plants expressing *LFY* under the *UBIQUITIN* promoter. (**A**) *LFY* transcript levels in the leaves of transgenic plants overexpressing *LFY.* The *LFY-A* coding region fused with an HA tag was driven by the maize *UBIQUITIN* promoter (*UBI:LFY- HA*). Five independent events were evaluated. Note the absence of expression of *LFY* in the negative transgenic sister lines (Neg.) and the wildtype Kronos control. *P* values are two-tail *t-* tests of each transgenic event (n = 4) relative to the combined Neg. and wildtype Kronos control (n = 8). * *P* = 0.05, ** *P* = 0.01, *** *P* = 0.001. Error bars are SEM. Raw data and statistics are available in data S4. (**B**) Floral defects in *UBI:LFY-HA* included reduced number or fused lodicules, fused anther filaments and extra stigmas. (**C**) Frequency of floral defects in 14 florets. Note the higher proportion of normal plants compared to the *lfy* mutants in Fig. S1B.

**Fig. S3.**
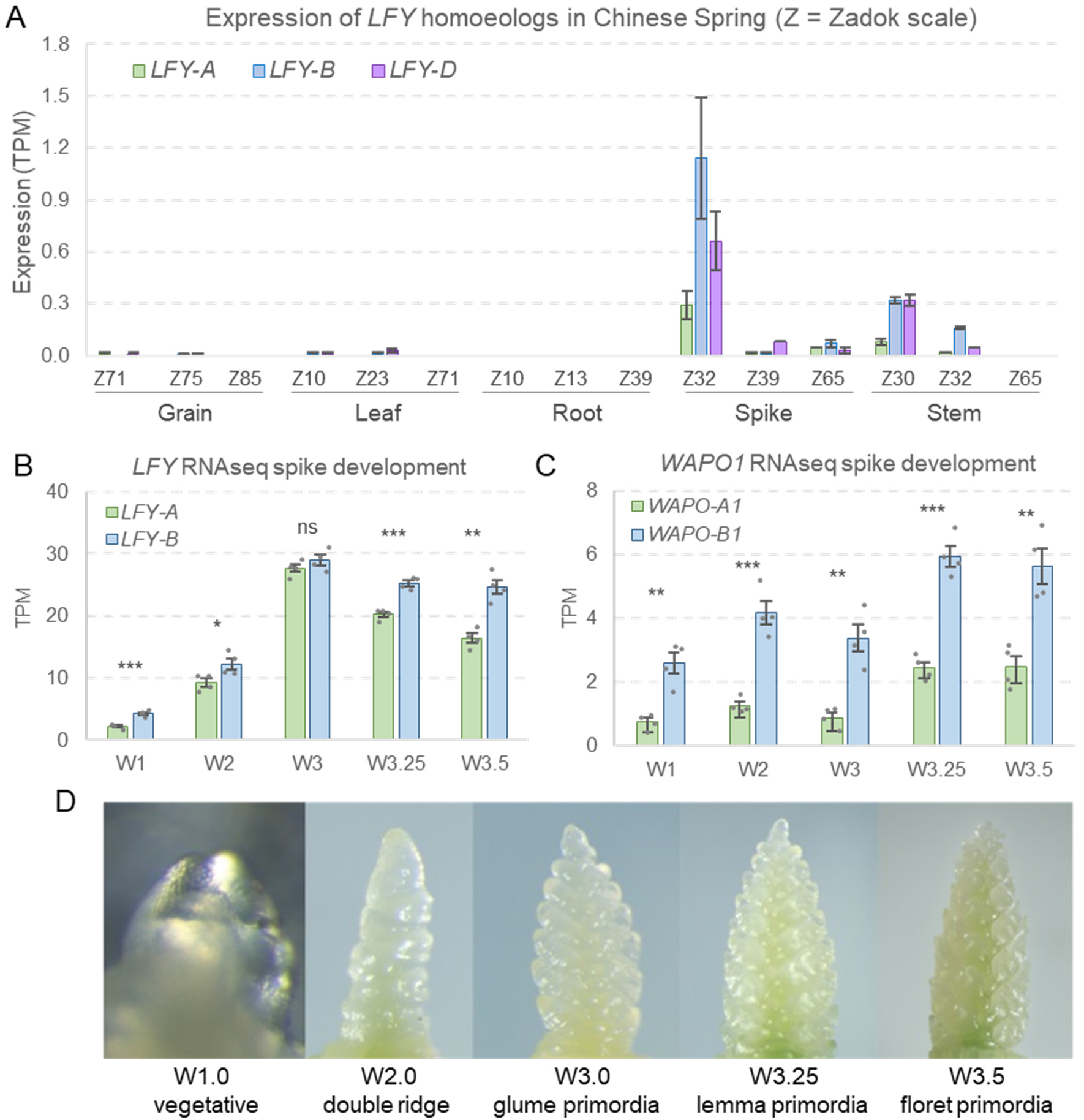
**Expression analysis of *LFY* and *WAPO1***. (**A**) *LFY* RNA-seq data for Chinese Spring across five tissues and three developmental stages (Choulet *et al*., 2014). (**B**) *LFY* and (**C**) *WAPO1* transcript levels during early stages of Kronos spike development (VanGessel *et al*., 2022). (**D**) Examples of developing spikes at the different spike development stages collected in the Kronos RNA-seq study (VanGessel *et al*., 2022). W numbers indicate values in the Waddington scale of wheat spike development (Waddington *et al*., 1983). TPM= transcripts per million. Averages are based on four biological replications per stage in each experiment and error bars are SEM. *P* values are from *t-*tests between homoeologs. ns= not significant, * = *P* < 0.05 and ** = *P* < 0.01. Raw data and statistics are available in data S7.

**Fig. S4.**
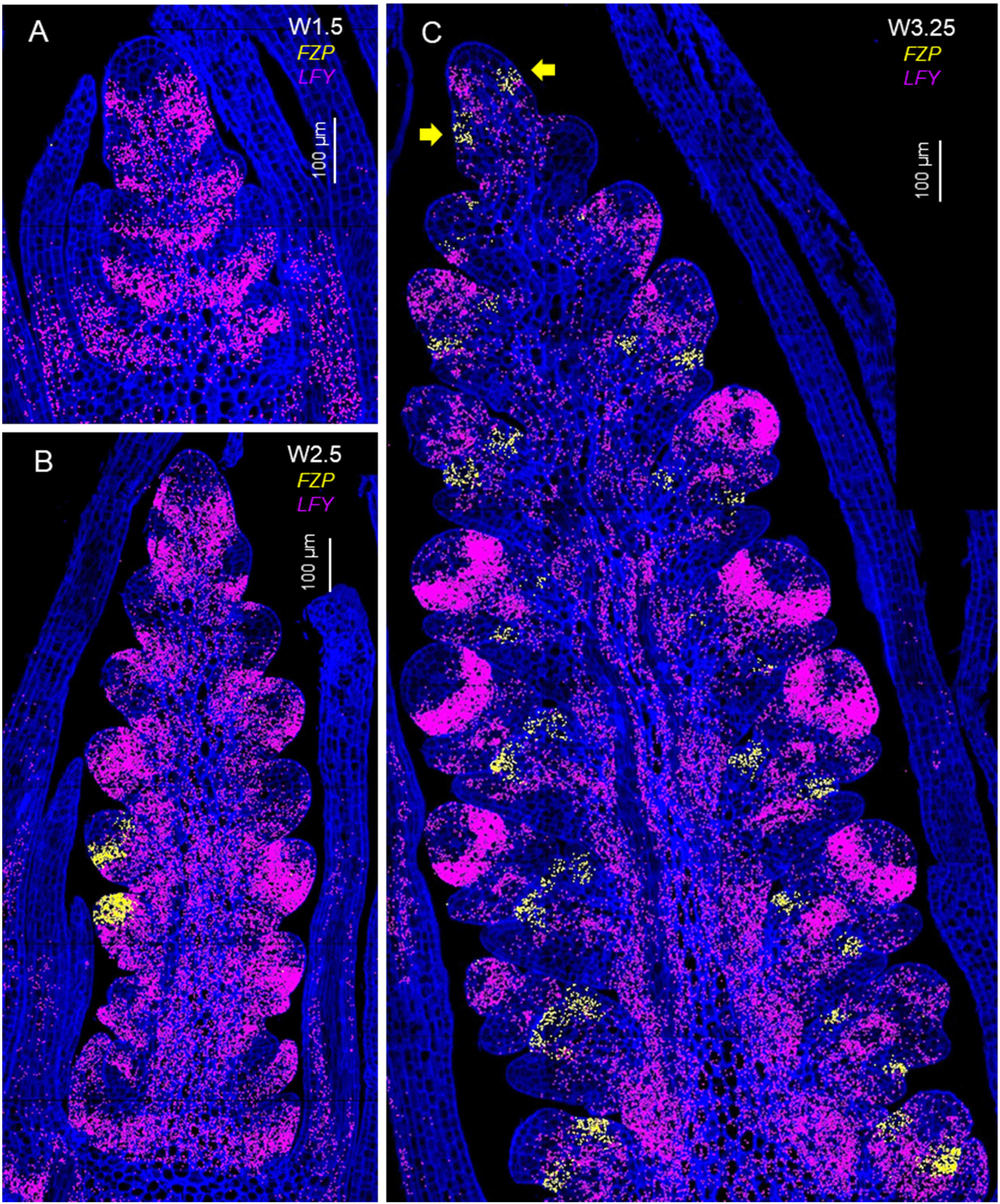
**Single-molecule fluorescence *in situ* hybridization (smFISH) for *FRIZZY PANICLE* (*FZP*).** *FZP* (*TraesCS2A02G116900*) is used as a marker for the initiation of spikelet development. *FZP* transcripts are indicated by yellow dots and *LFY* by purple dots. Cell walls stained with calcofluor are in dark blue. (**A**) Elongated shoot apical meristem transitioning to the reproductive stage (W1.5). (**B**) Late double ridge (DR, W2.5). (**C**) Lemma primordia (LP, W3.25). Yellow arrows indicate *FZP* expression in the youngest lateral meristems, suggesting a transition to glumes and the start of the IM transition to a terminal spikelet. Quantification of the *FZP* signal is available in data S8.

**Fig. S5.**
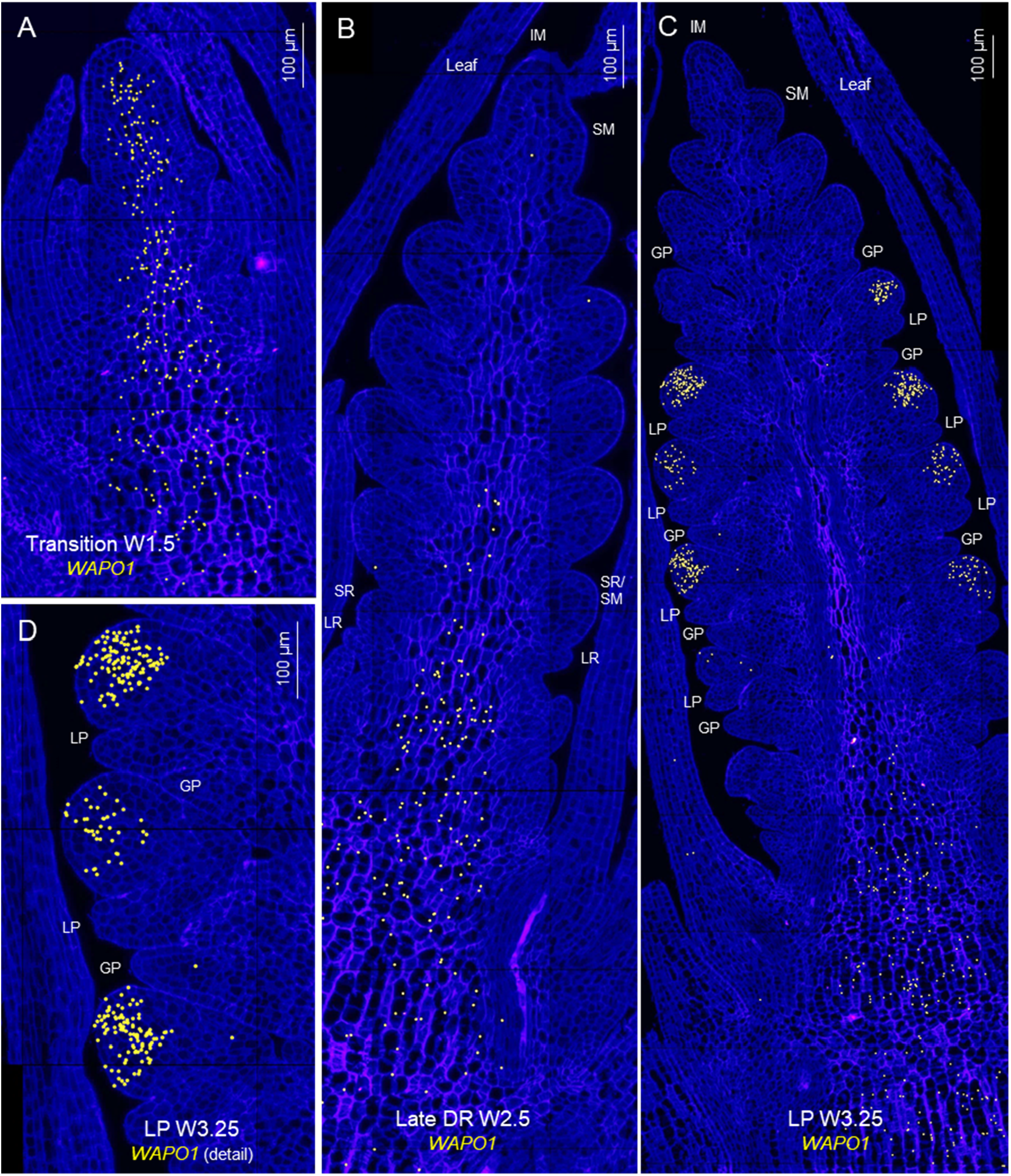
**Single-molecule fluorescence *in situ* hybridization (smFISH) for *WAPO1***. *WAPO1* (*TraesCS7B02G384000*) transcripts are indicated with yellow dots. Cell walls stained with calcofluor are presented in dark blue. (**A**) Elongated shoot apical meristem transitioning to the reproductive stage (W1.5). (**B**) Late double ridge (DR) stage (W2.5). (**C**) Lemma primordia (LP) stage (W3.25). (**D**) Detail of developing spikelet in C. LR = leaf ridge, SR = spikelet ridge, GP= glume primordium, LP= lemma primordium, IM = inflorescence meristem, SM= spikelet meristem (only one labelled). Scale bars are 100 μm.

**Fig. S6.**
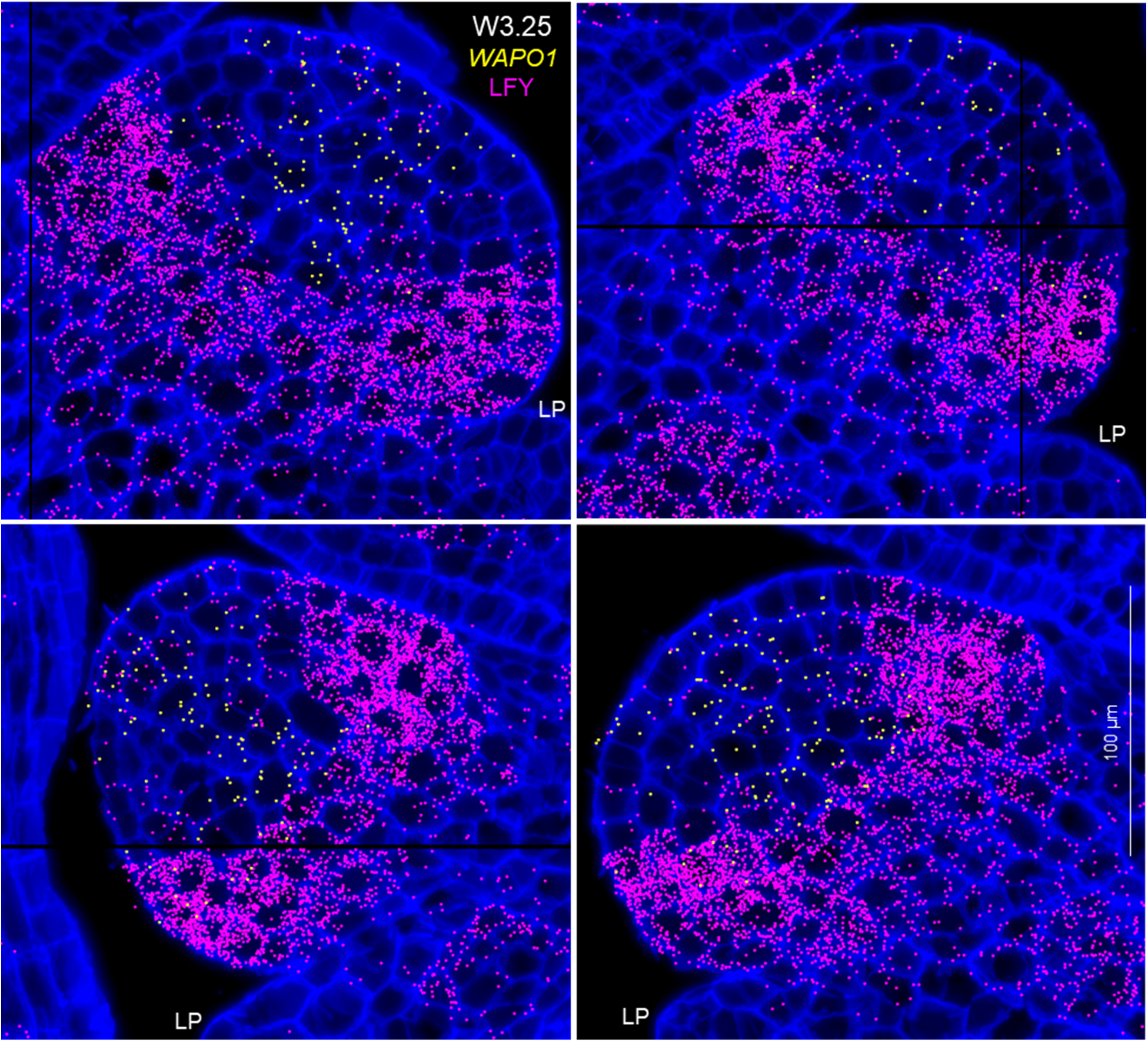
Single-molecule fluorescence *in situ* hybridization (smFISH) showing detail of the overlap between *LFY* and *WAPO1* in spikelet meristems. Spikelet meristems at the lemma primordia stage at W3.25. *WAPO1* transcripts are indicated with yellow dots and *LFY* transcript with purple dos. Cell walls stained with calcofluor are presented in dark blue. LP= lemma primordium. Scale bars are 100 μm.

**Fig. S7.**
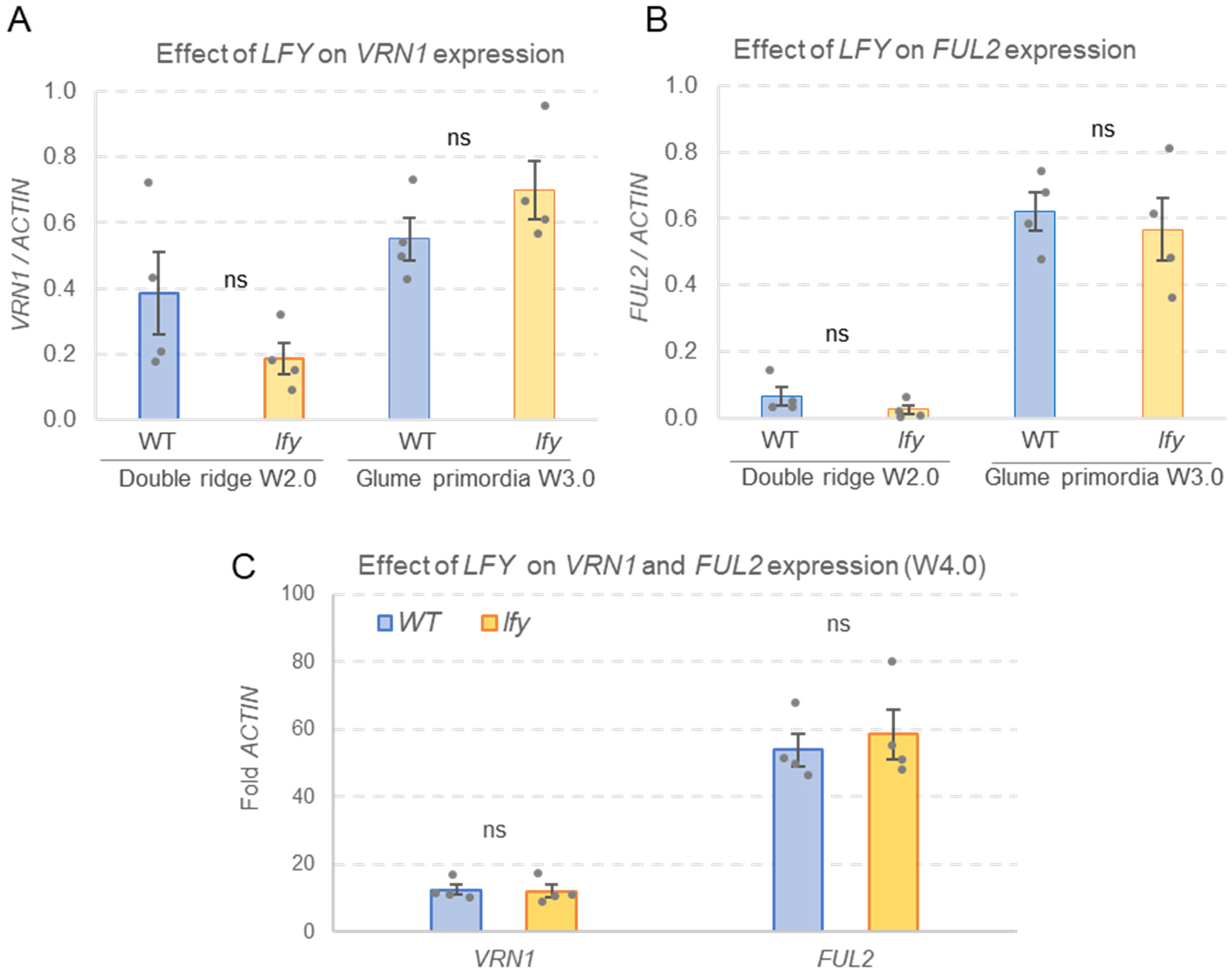
**Effect of *lfy* mutations on *VRN1* and *FUL2* expression**. Transcript levels were estimated by qRT-PCR relative to *ACTIN* endogenous control. (**A**) *VRN1* transcript levels at W.2.0 and W3.0 (**B**) *FUL2* transcript levels at W.2.0 and W3.0. (**C**) *VRN1* and *FUL2* transcript levels at W4.0. Bars are averages of four biological replicates and error bars are SEM. Each biological replicate represents one RNA extraction from a pool of four apices from Kronos wildtype (WT) and *lfy* mutant at the double ridge (W2.0), glume primordia (W3.0) or stamen primordium (4.0) stages of spike development. ns = not significant. Raw data and statistics are available in data S10.

**Fig. S8.**
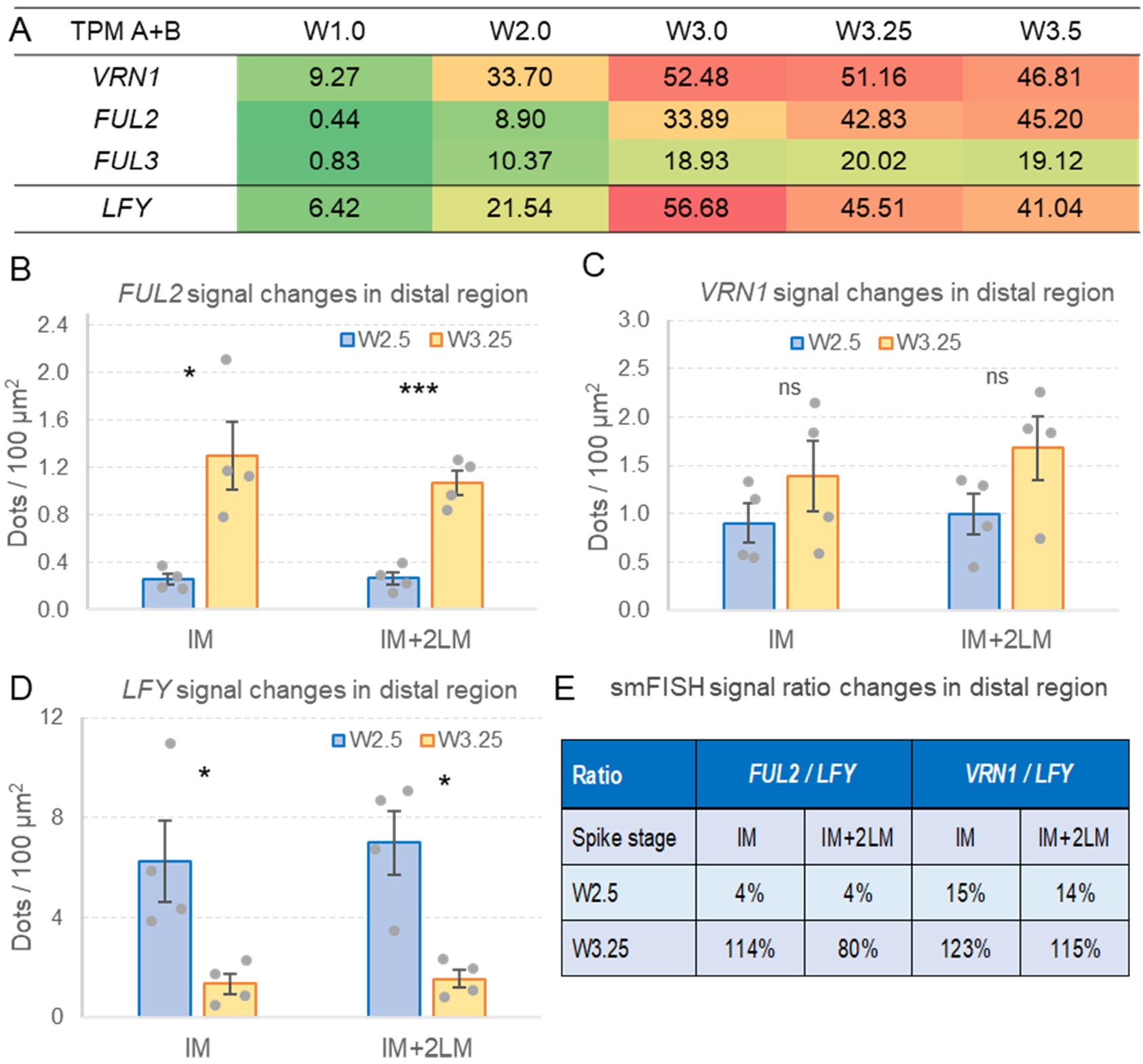
Changes in *VRN1*, *FUL2* and *LFY* between W2.5 and W3.25. (**A**) *VRN1*, *FUL2*, *FUL3* and *LFY* transcripts per million (TPM) from developing spikes at five stages (VanGessel *et al*., 2022). Values are the sum of A and B genome averages of four replications. Values for each genome are available in data S8. (**B-D**) Changes in hybridization signal density between late double ridge stage (W2.5) and lemma primordia stage (W3.25) at the inflorescence meristem (IM) and the IM plus the two youngest lateral meristems (IM+2LM). (**B**) *FUL2*, (**C**) *VRN1*, (**D**) *LFY.* (**E**) *FUL2/LFY* and *VRN1/LFY* ratios between hybridization signals at the IM and IM+2LM at W2.5 and W3.25. Values are averages of four sections at each stage (smFISH figures in data S8) and error bars are s.e.m. ns = not significant, * = *P* < 0.05 and *** = *P* < 0.001. Raw data and statistics are available in data S8.

**Fig. S9.**
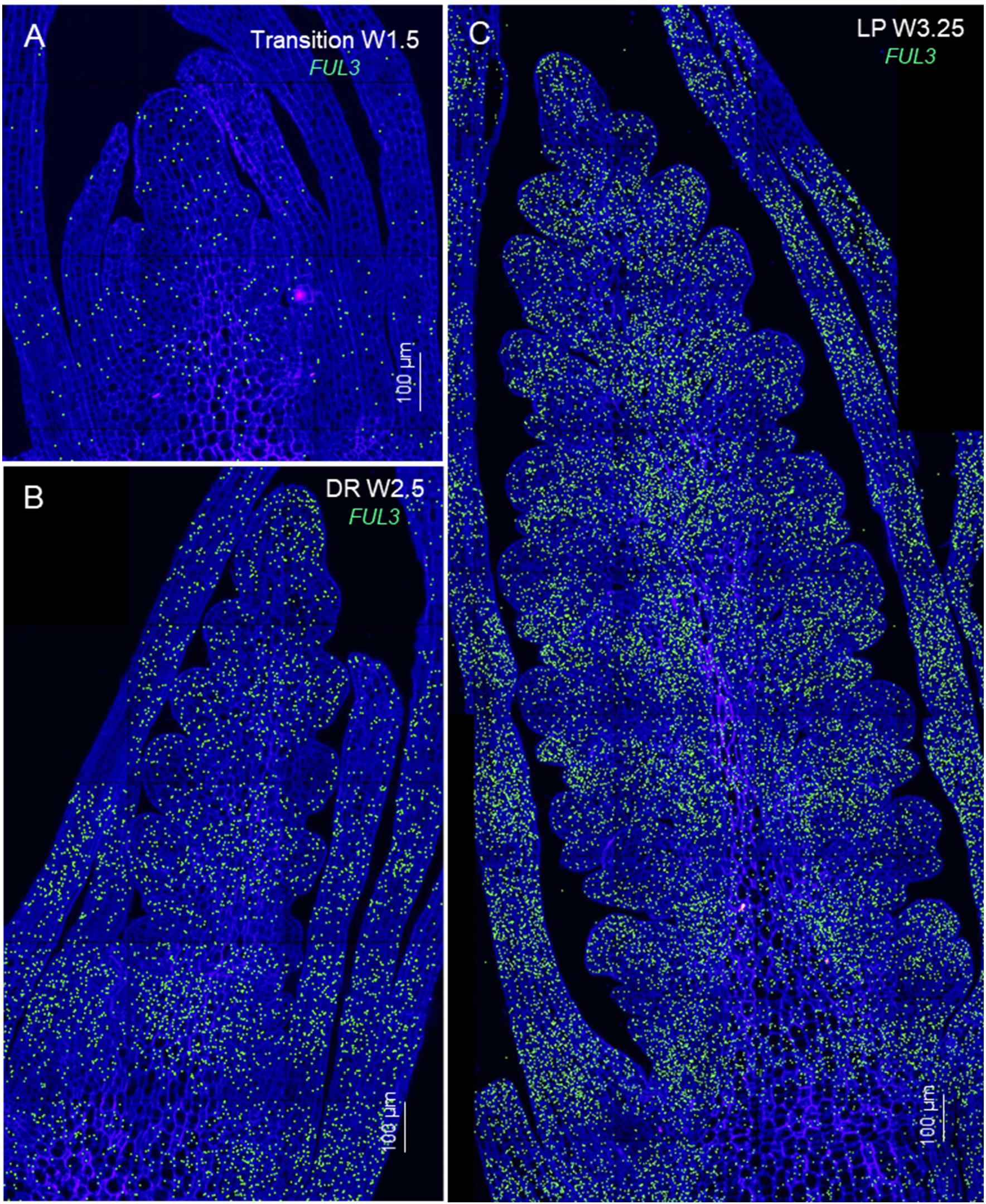
**Single-molecule fluorescence *in situ* hybridization (smFISH) for *FUL3***. Cell walls stained with calcofluor are presented in dark blue. (**A**) Elongated shoot apical meristem transitioning to the reproductive stage (W1.5). (**B**) Late double ridge stage (W2.5). (**C**) Lemma primordia stage (W3.25). Scale bars are 100 μm.

**Fig. S10.**
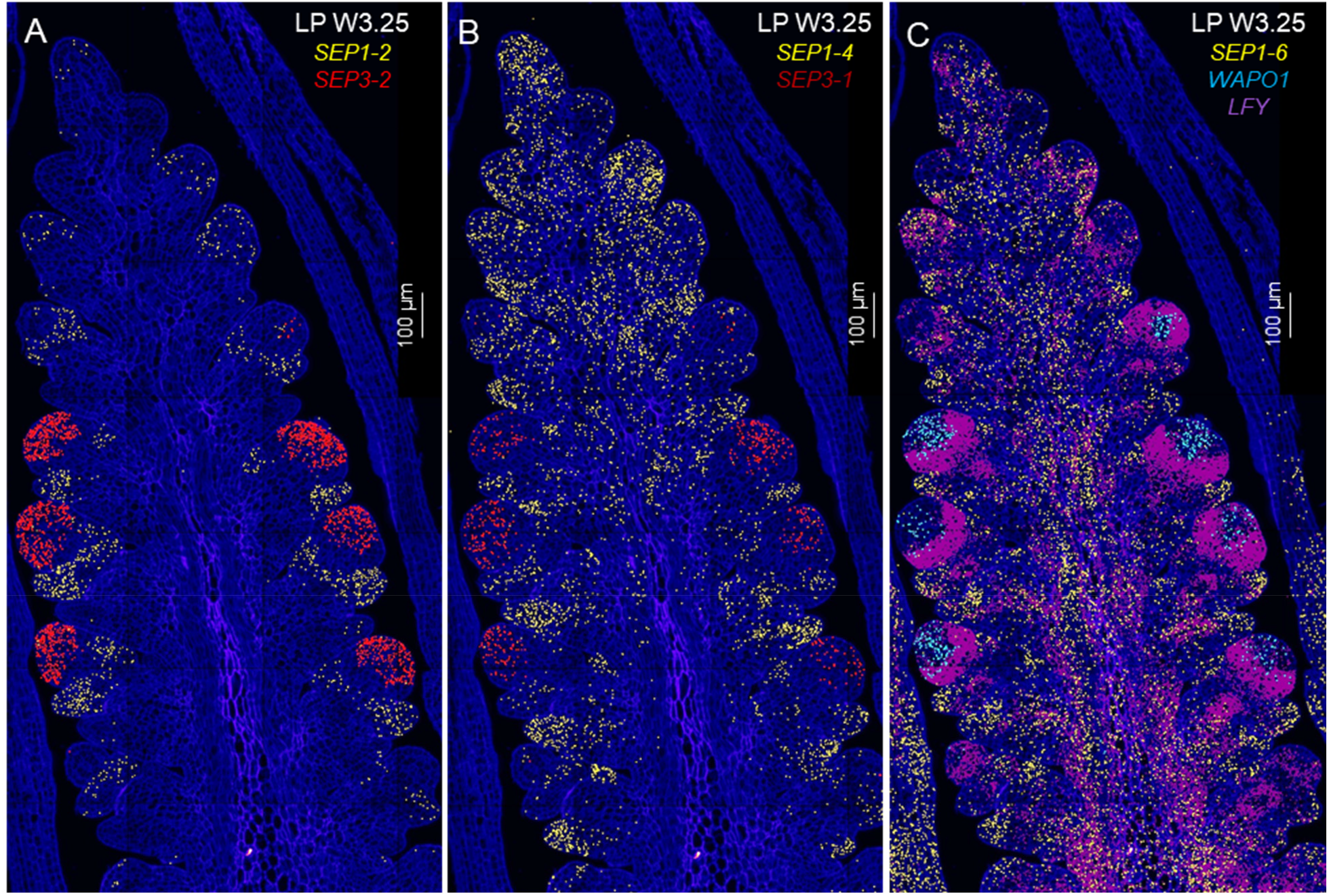
Single-molecule fluorescence *in situ* hybridization (smFISH) for *SEPALLATA* genes. Only the lemma primordia stage (W3.25) is presented. Cell walls stained with calcofluor are presented in dark blue. (**A**) *SEP1-2* (yellow) and *SEP3-2* (red). (**B**) *SEP1-4* (yellow) and *SEP3-1* (red). (**C**) *SEP1-6* (yellow), *WAPO1* (light blue) and *LFY* (violet). Scale bars are 100 μm. Gene identifications based on CS RefSeq v1.1 and rice orthologs are provided in data S9.

**Fig. S11.**
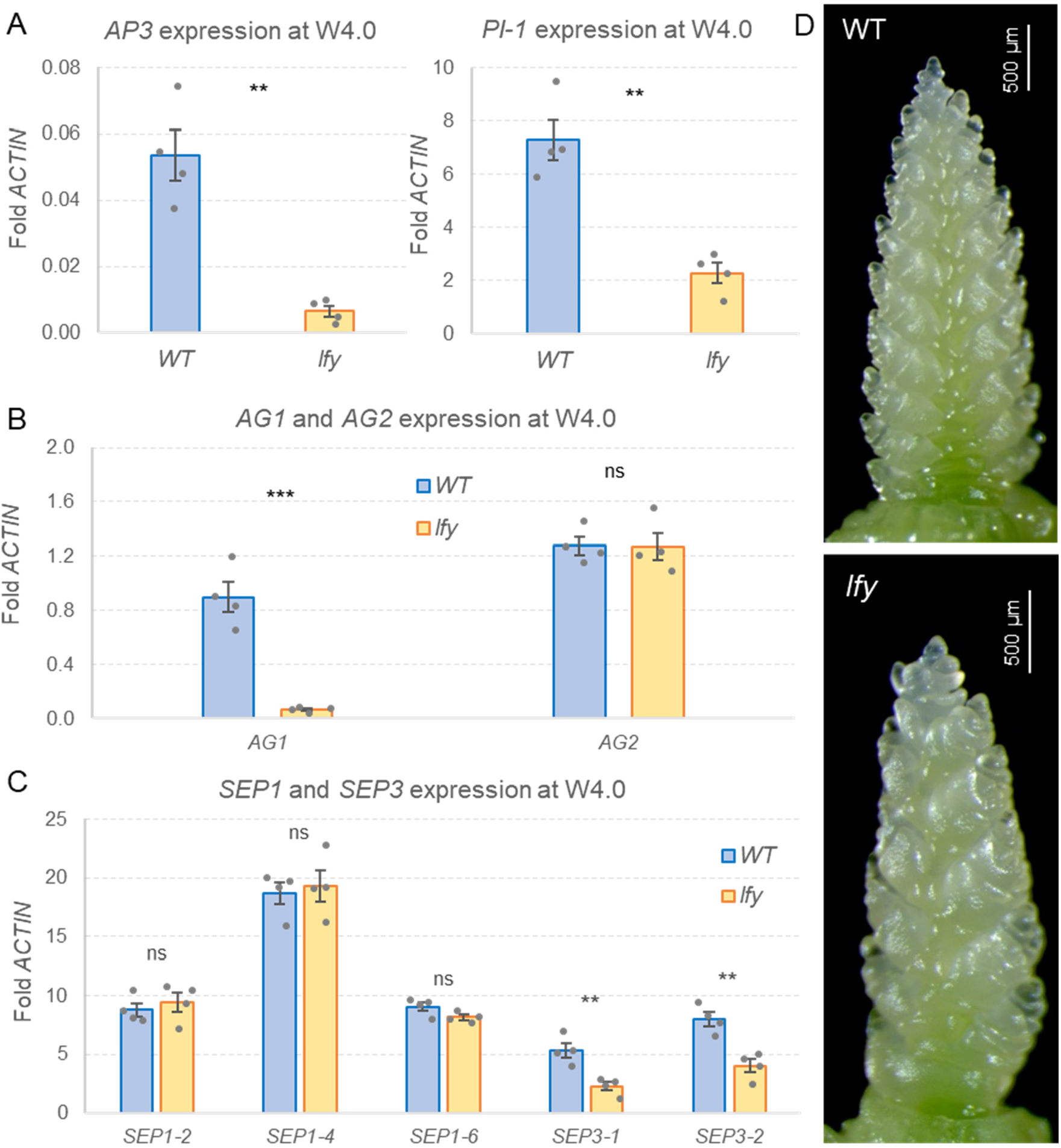
**Effect of *lfy* mutations on the expression of floral organ identity genes**. Expression of class-B, -C, and -E MADS-box genes was characterized in wheat developing spikes at the stamen primordia stage (W4.0) using qRT-PCR with *ACTIN* as endogenous control. (**A**) class-B genes *AP3-1* and *PI1*, (**B**) class-C genes *AG1* and *AG2*, (**C**) class-E genes *SEP1-2*, *SEP1-4*, *SEP1-6*, *SEP3-1*, *SEP3-2*. (**D**) Representative wheat developing spikes at W4.0 in WT Kronos and *lfy* mutants (note the reduced SNS). Bars are averages of four biological replicates and error bars are standard errors of the means (SEM). Each biological replicate represents one RNA extraction from a pool of four apices from Kronos wildtype (WT) and *lfy* mutant. ns = not significant, ** *P* < 0.01, *** *P* < 0.001. Raw data and statistics are available in data S11.

**Fig. S12.**
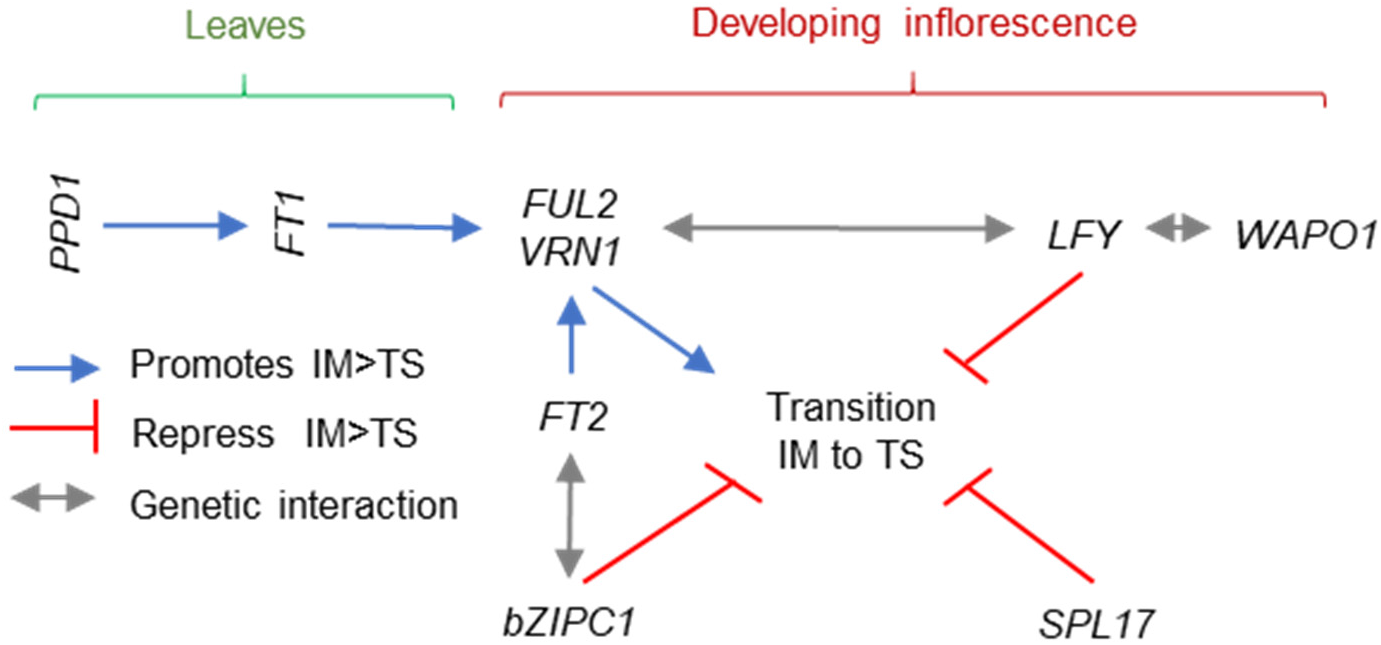
**Simplified working model of the regulation of SNS in wheat**. The negative effect of *bZIPC1* in the regulation of the transition between the inflorescence meristem to a terminal spikelet (IM>TS), and its interaction with *FT2* is based on (Glenn *et al*., 2023), the effect of *SPL17* on SNS in wheat is based on (Liu *et al*., 2023). The positive effects of *FT1* and *FT2* on the regulation of *VRN1* are based on (Li and Dubcovsky, 2008, Lv *et al*., 2014, Li *et al*., 2019, Shaw *et al*., 2019), and the positive effect of *PPD1* in the regulation of *FT1* and *FT2* on (Shaw *et al*., 2013).

## REFERENCES

1. Alvarez, M.A., Tranquilli, G., Lewis, S., Kippes, N. and Dubcovsky, J. (2016) Genetic and physical mapping of the earliness *per se* locus *Eps-A*^m^*1* in *Triticum monococcum* identifies *EARLY FLOWERING 3* (*ELF3*) as a candidate gene. Funct Integr Genomic, 16, 365–382.

2. Bomblies, K., Wang, R.L., Ambrose, B.A., Schmidt, R.J., Meeley, R.B. and Doebley, J. (2003) Duplicate *FLORICAULA/LEAFY* homologs *zfl1* and *zfl2* control inflorescence architecture and flower patterning in maize. Development, 130, 2385–2395.

3. Chae, E., Tan, Q.K.G., Hill, T.A. and Irish, V.F. (2008) An Arabidopsis F-box protein acts as a transcriptional co-factor to regulate floral development. Development, 135, 1235–1245.

4. Chen, A. and Dubcovsky, J. (2012) Wheat TILLING mutants show that the vernalization gene *VRN1* down-regulates the flowering repressor *VRN2* in leaves but is not essential for flowering. Plos Genet, 8, e1003134.

5. Choulet, F., Alberti, A., Theil, S., Glover, N., Barbe, V., Daron, J., Pingault, L., Sourdille, P., Couloux, A., Paux, E., Leroy, P., Mangenot, S., Guilhot, N., Le Gouis, J., Balfourier, F., Alaux, M., Jamilloux, V., Poulain, J., Durand, C., Bellec, A., Gaspin, C., Safar, J., Dolezel, J., Rogers, J., Vandepoele, K., Aury, J.M., Mayer, K., Berges, H., Quesneville, H., Wincker, P. and Feuillet, C. (2014) Structural and functional partitioning of bread wheat chromosome 3B. Science, 345, 1249721.

6. Coen, E.S., Romero, J.M., Doyle, S., Elliott, R., Murphy, G. and Carpenter, R. (1990) *Floricaula* - a homeotic gene required for flower development in *Antirrhinum majus*. Cell, 63, 1311–1322.

7. Debernardi, J.M., Greenwood, J.R., Jean Finnegan, E., Jernstedt, J. and Dubcovsky, J. (2020a) *APETALA 2*-like genes *AP2L2* and *Q* specify lemma identity and axillary floral meristem development in wheat. Plant J., 101, 171–187.

8. Debernardi, J.M., Tricoli, D.M., Ercoli, M.F., Hayta, S., Ronald, P., Palatnik, J.F. and Dubcovsky, J. (2020b) A GRF-GIF chimeric protein improves the regeneration efficiency of transgenic plants. Nat. Biotechnol., 38, 1274–1279.

9. FAOSTAT (2017) http://www.fao.org/faostat/en/#data: Food and Agriculture Organization (FAO) of the United Nations.

10. Ferrándiz, C., Gu, Q., Martienssen, R. and Yanofsky, M.F. (2000) Redundant regulation of meristem identity and plant architecture by *FRUITFULL*, APETALA1 and CAULIFLOWER. Development, 127, 725–734.

11. Frank, A.B. and Bauer, A. (1982) Effect of temperature and fertilizer-N on apex development in spring wheat. Agron. J., 74, 504–509.

12. Frank, A.B., Bauer, A. and Black, A.L. (1987) Effects of air-temperature and water-stress on apex development in spring wheat. Crop Sci., 27, 113–116.

13. Glenn, P., Woods, D.P., Zhang, J., Gabay, G., Odle, N. and Dubcovsky, J. (2023) Wheat bZIPC1 interacts with FT2 and contributes to the regulation of spikelet number per spike. Theor. Appl. Genet., 136, 237.

14. Glenn, P., Zhang, J., Brown-Guedira, G., DeWitt, N., Cook, J.P., Li, K., Akhunov, E. and Dubcovsky, J. (2022) Identification and characterization of a natural polymorphism in *FT-A2* associated with increased number of grains per spike in wheat. Theor Appl Genet, 135, 679–692.

15. Goslin, K., Zheng, B.B., Serrano-Mislata, A., Rae, L., Ryan, P.T., Kwasniewska, K., Thomson, B., O’Maoiléidigh, D.S., Madueño, F., Wellmer, F. and Graciet, E. (2017) Transcription factor interplay between LEAFY and APETALA1/CAULIFLOWER during floral initiation. Plant Physiol., 174, 1097–1109.

16. Honma, T. and Goto, K. (2000) The *Arabidopsis* floral homeotic gene *PSTILLATA* is regulated by discrete *cis*-elements responsive to induction and maintenance signals. Development, 127, 2021–2030.

17. Huala, E. and Sussex, I.M. (1992) Leafy interacts with floral homeotic genes to regulate Arabidopsis floral development. Plant Cell, 4, 901–913.

18. Ikeda-Kawakatsu, K., Maekawa, M., Izawa, T., Itoh, J.I. and Nagato, Y. (2012) *ABERRANT PANICLE ORGANIZATION 2/RFL*, the rice ortholog of Arabidopsis *LEAFY*, suppresses the transition from inflorescence meristem to floral meristem through interaction with *APO1*. Plant J., 69, 168–180.

19. Ikeda, K., Ito, M., NagasawaO, N., Kyozuka, J. and Nagato, Y. (2007) Rice *ABERRANT PANICLE ORGANIZATION 1*, encoding an F-box protein, regulates meristem fate. Plant J., 51, 1030–1040.

20. Ikeda, K., Nagasawa, N. and Nagato, Y. (2005) *ABERRANT PANICLE ORGANIZATION 1* temporally regulates meristem identity in rice. Dev. Biol., 282, 349–360.

21. Jin, R., Klasfeld, S., Zhu, Y., Garcia, M.F., Xiao, J., Han, S.K., Konkol, A. and Wagner, D. (2021) LEAFY is a pioneer transcription factor and licenses cell reprogramming to floral fate. Nat. Commun., 12, 626.

22. Komatsu, M., Chujo, A., Nagato, Y., Shimamoto, K. and Kyozuka, J. (2003) *FRIZZY PANICLE* is required to prevent the formation of axillary meristems and to establish floral meristem identity in rice spikelets. Development, 130, 3841–3850.

23. Krasileva, K.V., Vasquez-Gross, H.A., Howell, T., Bailey, P., Paraiso, F., Clissold, L., Simmonds, J., Ramirez-Gonzalez, R.H., Wang, X., Borrill, P., Fosker, C., Ayling, S., Phillips, A.L., Uauy, C. and Dubcovsky, J. (2017) Uncovering hidden variation in polyploid wheat. Proc. Natl. Acad. Sci. USA, 114, E913–E921.

24. Kuzay, S., Lin, H., Li, C., Chen, S., Woods, D., Zhang, J. and Dubcovsky, J. (2022) *WAPO- A1* is the causal gene of the 7AL QTL for spikelet number per spike in wheat. PLoS Genet., 18, e1009747.

25. Kuzay, S., Xu, Y., Zhang, J., Katz, A., Pearce, S., Su, Z., Fraser, M., Anderson, J.A., Brown-Guedira, G., DeWitt, N., Peters Haugrud, A., Faris, J.D., Akhunov, E., Bai, G. and Dubcovsky, J. (2019) Identification of a candidate gene for a QTL for spikelet number per spike on wheat chromosome arm 7AL by high-resolution genetic mapping. Theor Appl Genet, 132, 2689–2705.

26. Kyozuka, J., Konishi, S., Nemoto, K., Izawa, T. and Shimamoto, K. (1998) Down-regulation of *RFL*, the *FLO/LFY* homolog of rice, accompanied with panicle branch initiation. Proc. Natl. Acad. Sci. USA, 95, 1979–1982.

27. Lai, X.L., Blanc-Mathieu, R., GrandVuillemin, L., Huang, Y., Stigliani, A., Lucas, J., Thévenon, E., Loue-Manifel, J., Turchi, L., Daher, H., Brun-Hernandez, E., Vachon, G., Latrasse, D., Benhamed, M., Dumas, R., Zubieta, C. and Parcy, F. (2021) The LEAFY floral regulator displays pioneer transcription factor properties. Mol. Plant, 14, 829–837.

28. Lee, I., Wolfe, D.S., Nilsson, O. and Weigel, D. (1997) A *LEAFY* co-regulator encoded by *UNUSUAL FLORAL ORGANS*. Curr. Biol., 7, 95–104.

29. Lemmon, Z.H., Park, S.J., Jiang, K., Van Eck, J., Schatz, M.C. and Lippman, Z.B. (2016) The evolution of inflorescence diversity in the nightshades and heterochrony during meristem maturation. Genome Res., 26, 1676–1686.

30. Levin, J.Z. and Meyerowitz, E.M. (1995) *Ufo* - an Arabidopsis gene involved in both floral meristem and floral organ development. Plant Cell, 7, 529–548.

31. Li, C. and Dubcovsky, J. (2008) Wheat FT protein regulates *VRN1* transcription through interactions with FDL2. Plant J., 55, 543–554.

32. Li, C., Lin, H., Chen, A., Lau, M., Jernstedt, J. and Dubcovsky, J. (2019) Wheat *VRN1*, *FUL2* and *FUL3* play critical and redundant roles in spikelet development and spike determinacy. Development, 146, dev175398.

33. Li, C., Lin, H. and Dubcovsky, J. (2015) Factorial combinations of protein interactions generate a multiplicity of florigen activation complexes in wheat and barley. Plant J., 84, 70–82.

34. Liu, Y.Y., Chen, J., Yin, C.B., Wang, Z.Y., Wu, H., Shen, K.C., Zhang, Z.L., Kang, L.P., Xu, S., Bi, A.Y., Zhao, X.B., Xu, D.X., He, Z.H., Zhang, X.Y., Hao, C.Y., Wu, J.H., Gong, Y., Yu, X.C., Sun, Z.W., Ye, B.T., Liu, D.N., Zhang, L.L., Shen, L.P., Hao, Y.F., Ma, Y.Z., Lu, F. and Guo, Z.F. (2023) A high-resolution genotype-phenotype map identifies the *TaSPL17* controlling grain number and size in wheat. Genome Biol., 24, 196.

35. Lv, B., Nitcher, R., Han, X., Wang, S., Ni, F., Li, K., Pearce, S., Wu, J., Dubcovsky, J. and Fu, D. (2014) Characterization of *FLOWERING LOCUS T1* (*FT1*) gene in *Brachypodium* and wheat. PLoS One, 9, e94171.

36. Maas, E.V. and Grieve, C.M. (1990) Spike and leaf development in salt-stressed wheat. Crop Sci., 30, 1309–1313.

37. MacAlister, C.A., Park, S.J., Jiang, K., Marcel, F., Bendahmane, A., Izkovich, Y., Eshed, Y. and Lippman, Z.B. (2012) Synchronization of the flowering transition by the tomato *TERMINATING FLOWER* gene. Nat. Genet., 44, 1393–1398.

38. Maizel, A., Busch, M.A., Tanahashi, T., Perkovic, J., Kato, M., Hasebe, M. and Weigel, D. (2005) The floral regulator LEAFY evolves by substitutions in the DNA binding domain. Science, 308, 260–263.

39. Miao, Y.L., Xun, Q., Taji, T., Tanaka, K., Yasuno, N., Ding, C.Q. and Kyozuka, J. (2022) *ABERRANT PANICLE ORGANIZATION2* controls multiple steps in panicle formation through common direct-target genes. Plant Physiol., 189, 2210–2226.

40. Molinero-Rosales, N., Jamilena, M., Zurita, S., Gómez, P., Capel, J. and Lozano, R. (1999) *FALSIFLORA*, the tomato orthologue of *FLORICAULA* and *LEAFY*, controls flowering time and floral meristem identity. Plant J., 20, 685–693.

41. Pajoro, A., Madrigal, P., Muino, J.M., Matus, J.T., Jin, J., Mecchia, M.A., Debernardi, J.M., Palatnik, J.F., Balazadeh, S., Arif, M., O’Maoileidigh, D.S., Wellmer, F., Krajewski, P., Riechmann, J.L., Angenent, G.C. and Kaufmann, K. (2014) Dynamics of chromatin accessibility and gene regulation by MADS-domain transcription factors in flower development. Genome Biol., 15, R41.

42. Parcy, F., Nilsson, O., Busch, M.A., Lee, I. and Weigel, D. (1998) A genetic framework for floral patterning. Nature, 395, 561–566.

43. Preston, J.C., Christensen, A., Malcomber, S.T. and Kellogg, E.A. (2009) MADS-box gene expression and implications for developmental origins of the grass spikelet. Amer. Jour. Bot., 96, 1419–1429.

44. Ream, T.S., Woods, D.P., Schwartz, C.J., Sanabria, C.P., Mahoy, J.A., Walters, E.M., Kaeppler, H.F. and Amasino, R.M. (2014) Interaction of photoperiod and vernalization determines flowering time of *Brachypodium distachyon*. Plant Physiol., 164, 694–709.

45. Rieu, P., Arnoux-Courseaux, M., Tichtinsky, G. and Parcy, F. (2023a) Thinking outside the F-box: how UFO controls angiosperm development. New Phytol., 240, 945–959.

46. Rieu, P., Turchi, L., Thevenon, E., Zarkadas, E., Nanao, M., Chahtane, H., Tichtinsky, G., Lucas, J., Blanc-Mathieu, R., Zubieta, C., Schoehn, G. and Parcy, F. (2023b) The F-box protein UFO controls flower development by redirecting the master transcription factor LEAFY to new cis-elements. Nat. Plants, 9, 315–329.

47. Schultz, E.A. and Haughn, G.W. (1991) *LEAFY*, a homeotic gene that regulates inflorescence development in Arabidopsis. Plant Cell, 3, 771–781.

48. Selva, C., Shirley, N.J., Houston, K., Whitford, R., Baumann, U., Li, G. and Tucker, M.R. (2021) *HvLEAFY* controls the early stages of floral organ specification and inhibits the formation of multiple ovaries in barley. Plant J., 108, 509–527.

49. Shaw, L.M., Lyu, B., Turner, R., Li, C., Chen, F., Han, X., Fu, D. and Dubcovsky, J. (2019) *FLOWERING LOCUS T2* regulates spike development and fertility in temperate cereals. J. Exp. Bot., 70, 193–204.

50. Shaw, L.M., Turner, A.S., Herry, L., Griffiths, S. and Laurie, D.A. (2013) Mutant alleles of *Photoperiod-1* in wheat (*Triticum aestivum* L.) that confer a late flowering phenotype in long days. PLoS One, 8, e79459.

51. Shitsukawa, N., Takagishi, A., Ikari, C., Takumi, S. and Murai, K. (2006) WFL, a wheat FLORICAULA/LEAFY ortholog, is associated with spikelet formation as lateral branch of the inflorescence meristem. Genes & Genetic Systems, 81, 13–20.

52. Souer, E., van der Krol, A., Kloos, D., Spelt, C., Bliek, M., Mol, J. and Koes, R. (1998) Genetic control of branching pattern and floral identity during inflorescence development. Development, 125, 733–742.

53. VanGessel, C., Hamilton, J., Tabbita, F., Dubcovsky, J. and Pearce, S. (2022) Transcriptional signatures of wheat inflorescence development. Sci Rep-Uk, 12, 17224.

54. Waddington, S.R., Cartwright, P.M. and Wall, P.C. (1983) A quantitative scale of spike initial and pistil development in barley and wheat. Ann. Bot., 51, 119–130.

55. Wagner, D., Sablowski, R.W.M. and Meyerowitz, E.M. (1999) Transcriptional activation of *APETALA1* by LEAFY. Science, 285, 582–584.

56. Weigel, D., Alvarez, J., Smyth, D.R., Yanofsky, M.F. and Meyerowitz, E.M. (1992) Leafy controls floral meristem identity in Arabidopsis. Cell, 69, 843–859.

57. Weigel, D. and Nilsson, O. (1995) A developmental switch sufficient for flower Initiation in diverse plants. Nature, 377, 495–500.

58. William, D.A., Su, Y.H., Smith, M.R., Lu, M., Baldwin, D.A. and Wagner, D. (2004) Genomic identification of direct target genes of LEAFY. Proc. Natl. Acad. Sci. USA, 101, 1775–1780.

59. Winter, C.M., Yamaguchi, N., Wu, M.F. and Wagner, D. (2015) Transcriptional programs regulated by both LEAFY and APETALA1 at the time of flower formation. Physiol. Plantarum, 155, 55–73.

60. Yamaguchi, N. (2021) LEAFY, a pioneer transcription factor in plants: A mini-review. Front. Plant. Sci., 12.

61. Yoshida, H. (2012) Is the lodicule a petal: Molecular evidence? Plant Sci., 184, 121–128.

62. Zhang, J., Li, C., Zhang, W., Zhang, X., Mo, Y., Tranquilli, G.E., Vanzetti, L.S. and Dubcovsky, J. (2023) Wheat plant height locus *RHT25* encodes a PLATZ transcription factor that interacts with DELLA (RHT1). Proc. Natl. Acad. Sci. USA, 120, e2300203120.

63. Zhang, J.L., Gizaw, S.A., Bossolini, E., Hegarty, J., Howell, T., Carter, A.H., Akhunov, E. and Dubcovsky, J. (2018) Identification and validation of QTL for grain yield and plant water status under contrasting water treatments in fall-sown spring wheats. Theor. Appl. Genet., 131, 1741–1759.

64. Zhong, J., van Esse, G.W., Bi, X., Lan, T., Walla, A., Sang, Q., Franzen, R. and von Korff, M. (2021) *INTERMEDIUM-M* encodes an HvAP2L-H5 ortholog and is required for inflorescence indeterminacy and spikelet determinacy in barley. P Natl Acad Sci USA, 118, e2011779118.

## References for Supplemental Figures

65. Choulet, F., Alberti, A., Theil, S., Glover, N., Barbe, V., Daron, J., Pingault, L., Sourdille, P., Couloux, A., Paux, E., Leroy, P., Mangenot, S., Guilhot, N., Le Gouis, J., Balfourier, F., Alaux, M., Jamilloux, V., Poulain, J., Durand, C., Bellec, A., Gaspin, C., Safar, J., Dolezel, J., Rogers, J., Vandepoele, K., Aury, J.M., Mayer, K., Berges, H., Quesneville, H., Wincker, P. and Feuillet, C. (2014) Structural and functional partitioning of bread wheat chromosome 3B. Science, 345, 1249721.

66. Glenn, P., Woods, D.P., Zhang, J., Gabay, G., Odle, N. and Dubcovsky, J. (2023) Wheat bZIPC1 interacts with FT2 and contributes to the regulation of spikelet number per spike. Theor. Appl. Genet., 136, 237.

67. Li, C. and Dubcovsky, J. (2008) Wheat FT protein regulates *VRN1* transcription through interactions with FDL2. Plant J., 55, 543–554.

68. Li, C., Lin, H., Chen, A., Lau, M., Jernstedt, J. and Dubcovsky, J. (2019) Wheat *VRN1*, *FUL2* and *FUL3* play critical and redundant roles in spikelet development and spike determinacy. Development, 146, dev175398.

69. Liu, Y.Y., Chen, J., Yin, C.B., Wang, Z.Y., Wu, H., Shen, K.C., Zhang, Z.L., Kang, L.P., Xu, S., Bi, A.Y., Zhao, X.B., Xu, D.X., He, Z.H., Zhang, X.Y., Hao, C.Y., Wu, J.H., Gong, Y., Yu, X.C., Sun, Z.W., Ye, B.T., Liu, D.N., Zhang, L.L., Shen, L.P., Hao, Y.F., Ma, Y.Z., Lu, F. and Guo, Z.F. (2023) A high-resolution genotype-phenotype map identifies the *TaSPL17* controlling grain number and size in wheat. Genome Biol., 24, 196.

70. Lv, B., Nitcher, R., Han, X., Wang, S., Ni, F., Li, K., Pearce, S., Wu, J., Dubcovsky, J. and Fu, D. (2014) Characterization of *FLOWERING LOCUS T1* (*FT1*) gene in *Brachypodium* and wheat. PLoS One, 9, e94171.

71. Shaw, L.M., Lyu, B., Turner, R., Li, C., Chen, F., Han, X., Fu, D. and Dubcovsky, J. (2019) *FLOWERING LOCUS T2* regulates spike development and fertility in temperate cereals. J. Exp. Bot., 70, 193–204.

72. Shaw, L.M., Turner, A.S., Herry, L., Griffiths, S. and Laurie, D.A. (2013) Mutant alleles of *Photoperiod-1* in wheat (*Triticum aestivum* L.) that confer a late flowering phenotype in long days. PLoS One, 8, e79459.

73. VanGessel, C., Hamilton, J., Tabbita, F., Dubcovsky, J. and Pearce, S. (2022) Transcriptional signatures of wheat inflorescence development. Sci Rep-Uk, 12, 17224.

74. Waddington, S.R., Cartwright, P.M. and Wall, P.C. (1983) A quantitative scale of spike initial and pistil development in barley and wheat. Ann. Bot., 51, 119–130.

